# Dynamics of the α2-chimaerin–Rac1 interface in Duane retraction syndrome and computational design of candidate probes

**DOI:** 10.64898/2026.07.29.741529

**Authors:** Anwesha Shah, Yash Bhargava

**Author notes:** Corresponding author: Yash Bhargava.

## Abstract

Duane retraction syndrome (DRS) is an eye movement disorder caused by gain-of-function mutations in CHN1, which encodes α2-chimaerin. Mutations in CHN1 cause excessive Rac1 suppression during axon guidance, but the dynamics of the α2-chimaerin–Rac1 interaction remain largely uncharacterized. We simulated the CHN1– Rac1 complex with all-atom molecular dynamics to characterize this interface. An Ile420–Arg445 hydrophobic ridge is the persistent core, while the 304–310 patch carrying the catalytic arginine finger scores lower and varies more between replicates. We screened the BioGRID partners of CHN1 with AlphaFold3 and ipSAE; only Rac1, NCK1, and NCK2 gave high-confidence predicted complexes. Rac1 and NCK1 stayed bound in every replicate, and NCK2 in two of three. The MM-GBSA interface ΔG for Rac1 was −20 ± 10 kcal/mol. We also explored how the surface could be probed by small molecules and peptides. To identify candidate chemical probes, we sampled natural products from COCONUT, refined top hits through CReM fragment growth, and generated peptides with BoltzGen. Prioritized ligands were evaluated with molecular dynamics and MM-PBSA endpoint free-energy calculations. Across 6 µs of cumulative simulation, the contact network defining the CHN1-Rac1 interface matched ligand and peptide engagement. These results define the key residues of the CHN1–Rac1 interface and identify candidate chemical probes for testing their role experimentally.

## Introduction

Duane Retraction Syndrome (DRS) is a rare eye movement congenital condition in which one or both eyes cannot move normally from side to side, often causing the eye to pull backward into the socket and the eyelid opening to narrow during attempted movement (Kekunnaya & Negalur, 2017). DRS is clinically divided into three main subtypes based on the pattern of movement limitation: Type I (limited abduction), Type II (limited adduction), and Type III (limitation of both) (Huber, 1974). Familial DRS attributable to *CHN1* mutations, designated DURS2, is typically bilateral (Chan et al., 2011; Miyake et al., 2008). DRS is within a broader class of congenital cranial dysinnervation disorders, in which disruptions in axon guidance and growth cone navigation during embryogenesis lead to persistent defects in neural circuitry (Angelini et al., 2021; Chan et al., 2011; Whitman & Engle, 2017).

Current management for DRS is almost entirely surgical and focuses on improving eye alignment or compensatory head posture. Horizontal rectus recession can improve alignment in primary gaze and reduce abnormal movements such as upshoots or downshoots but does not restore normal ocular motility or binocular coordination (Sheth et al., 2021). Prismatic glasses can correct eye misalignment and reduce head turning without surgery, but they suit only selected patients: in one study only 12 of 638 patients with DRS met the criteria for prism management, and although the misalignment in primary position was corrected in all 12, the abnormal head posture cleared completely in only 5 (Aygit et al., 2017). Since these interventions act at the level of the muscles, they cannot influence the primary developmental and molecular defect in CHN1-associated DRS or be applied during the embryonic window while ocular motor axons are forming their connections.

Genetic studies of familial DRS have identified heterozygous missense mutations in the *CHN1* gene as a recurrent cause of autosomal dominant disease (Chan et al., 2011; Miyake et al., 2008). *CHN1* encodes α2-chimaerin, a neuronally enriched Rac GTPase-activating protein (RacGAP) that regulates cytoskeletal dynamics during neural development by modulating Rac1 signalling (Hall & Lalli, 2010; Iwasato et al., 2007). Under normal conditions, α2-chimaerin exists in an autoinhibited state in the cytosol and is recruited to the membrane on binding diacylglycerol, where it inactivates Rac1 through GTP hydrolysis (Colón-González et al., 2008). Disease-associated mutations destabilize this autoinhibited conformation, leading to increased membrane association and excessive RacGAP activity (Colón-González et al., 2008; Miyake et al., 2008). This results in inappropriate suppression of Rac1 signalling during critical stages of axon pathfinding (Ferrario et al., 2012). Experimental models overexpressing mutant CHN1 demonstrate disrupted ocular motor axon trajectories and patterns of misinnervation that closely match human DRS, providing strong evidence that dysregulated α2-chimaerin–Rac1 signalling drives disease pathogenesis (Miyake et al., 2008).

Although CHN1-associated DRS has a defined molecular basis, the dynamics of the α2-chimaerin–Rac1 complex remain unknown. Protein–protein interfaces like this are often flat and flexible, so a static structure cannot fully explain their behavior. Molecular dynamics (MD) shows which contacts remain stable, how the proteins move, and how binding changes their motion. Alanine-scanning studies introduced the idea of interface “hot spots,” or residues that contribute most of the binding strength (Bogan & Thorn, 1998; Clackson & Wells, 1995). Integrated molecular-dynamics workflows are now a standard route to interrogating such surfaces (Khademi Dehkordi et al., 2024; Naik et al., 2024; Rehman et al., 2023). Additionally, diffusion-based generative models now enable the design of cyclic peptides capable of engaging complex protein surfaces with high affinity (Rettie et al., 2025). These approaches provide a way to identify chemical probes that can experimentally perturb the α2-chimaerin– Rac1 interface and reveal which interactions are most important.

Using the workflow in Fig. 1B, we relaxed a complex by MD, defined the interface from trajectory-derived contacts, tested whether Rac1 is a genuine partner against the reported CHN1 interactome, and asked how Rac1 binding changes the collective motion of CHN1 itself. We then used the resulting contact map to design small molecules and cyclic peptides against it.

**Figure 1.**
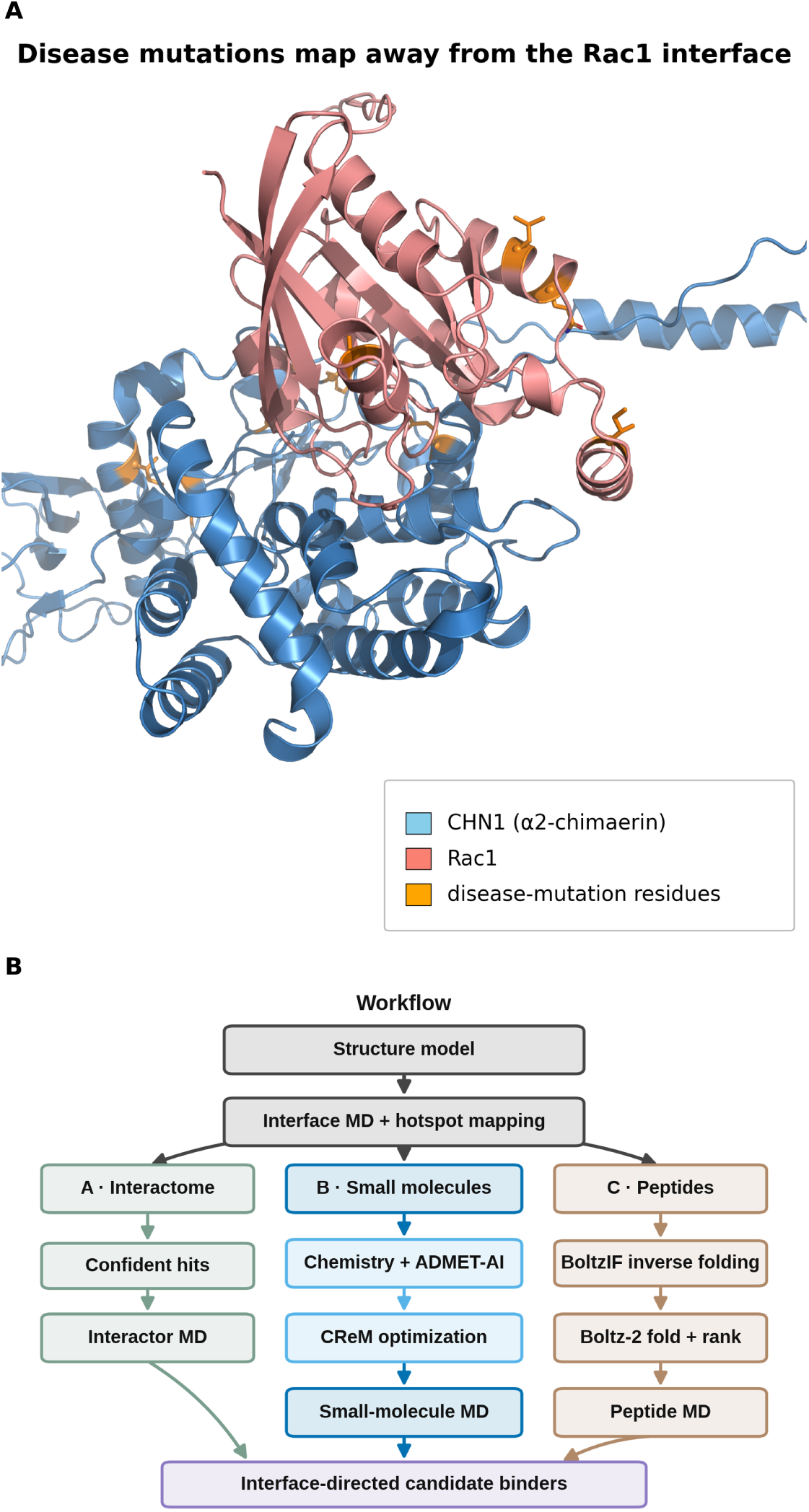
The CHN1–Rac1 model and the workflow built on it. **(A)** The hybrid CHN1–Rac1 assembly, combining the experimental CHN1 fold (PDB 3CXL; structural Zn^2+^ retained) with the AlphaFold3-predicted binding orientation and the reconstructed 152–202 loop (see Methods). CHN1 is blue, Rac1 salmon. The nine mapped disease-associated substitutions, which fall on eight residues because Pro252 carries two, are drawn as orange side-chain sticks with a Cα sphere; all lie distal to the Rac1 interface, consistent with an allosteric mechanism. **(B)** Overview of the workflow: structure assembly and MD-based interface-contact mapping feed three arms, an AlphaFold3/ipSAE interactor survey with interactor MD, COCONUT sampling with CReM optimization and ADMET triage, and BoltzGen cyclic-peptide design, each validated by MD.

## Materials and methods

### Structural modeling and definition of the CHN1–Rac1 interface

CHN1 was modeled from UniProt P15882-1 (residues 15–459, the span resolved in the crystal structure) and Rac1 from P63000 (residues 1–192). Wild-type CHN1 was used because CHN1-associated DRS mutations raise activity by destabilizing autoinhibited conformations rather than by substituting Rac1-binding surface residues (Miyake et al., 2008). Nine reported CHN1 substitutions (Leu20Phe, Ile126Met, Tyr143His, Pro141Leu, Ala223Val, Gly228Ser, Pro252Gln, Pro252Ser, Glu313Lys) were mapped onto the model and all lie more than 4 Å from the predicted interface (Biler et al., 2017; Chan et al., 2011; Miyake et al., 2008; see Figure 1B and Table S1).

An initial CHN1–Rac1 complex was generated with the AlphaFold3 web server at default settings (Abramson et al., 2024), scoring pTM 0.70 and ipTM 0.74. Rac1 is modeled and simulated apo throughout and Rac1 switch I (residues 26–45) and switch II (59–76) were compared with PDB 1MH1, 1G4U and 1HH4 by superposing and measuring each switch separately. The CHN1 chain was rebuilt on the experimental crystal structure (PDB 3CXL; 2.60 Å; Shen et al., 2008). We superposed the ordered regions of the AlphaFold3 model on 3CXL, kept the 3CXL coordinates as the scaffold, and built the two unresolved segments (residues 152–160 and 168–202) from the AlphaFold3 model as one aligned span covering residues 148–203. We validated the built and MD-relaxed model using Ramachandran analysis. Residues were classified using the MDAnalysis reference density and grouped by whether they came from the original structure or were newly built (Supplementary Fig. S1). Both structural zinc ions were retained and the crystallographic UNK heteroatom removed. The hybrid model combines the experimentally resolved and predicted structures to get a full length construct for modeling (Fig. 1A). As an independent check on the interface region, 3CXL was superposed on the separately determined 1.8 Å structure of the same RhoGAP domain (PDB 2OSA; Walker et al., 2007) over their 196 common Cα atoms. The predicted chain orientation was compared with an experimental GAP–Rac1 complex. Our Rac1 chain was superposed on that of PDB 1G4U, *Salmonella* SptP bound to Rac1 with GDP, Mg^2+^ and AlF_3_ (Stebbins & Galán, 2000). The CHN1 arginine finger was then compared with SptP Arg209, measured guanidinium CZ to CZ and from CZ to the AlF_3_ centroid.

The hybrid model was relaxed by MD for 200 ns with frames analyzed under the CHARMM36 protocol detailed below. Trajectories were corrected for periodic-boundary artifacts and aligned on the protein backbone before analysis.

The interface was defined from trajectory-derived contacts using MDAnalysis (Gowers et al., 2016; Michaud-Agrawal et al., 2011), with CHN1 as chain A and Rac1 as chain B. A contact was recorded when opposing-chain heavy atoms fell within 4.5 Å in a frame.

For each CHN1 residue *i* an interface-contact score on a 0–100 scale was computed as

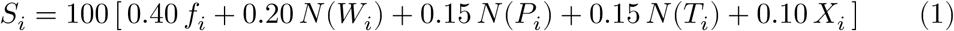

where *f_i_* is the fraction of frames in which residue *i* contacts Rac1; *W_i_* is the sum, over Rac1 partner residues, of their pairwise contact occupancies; *P_i_* is the number of Rac1 partners contacted in at least 20 % of frames; *T_i_* is the interaction-type term of Eq. (2); *X_i_* is the proximity term of Eq. (3); and N(·) denotes division by the maximum value of that quantity over all CHN1 residues within the same trajectory.

The interaction-type term sums the four contact classes with fixed weights,

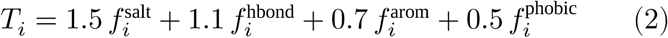

each *f* being the fraction of frames in which the residue makes a contact of that class. A salt bridge was counted when a charged side-chain nitrogen (Lys NZ; Arg NE, NH1, NH2; His ND1, NE2) and a charged side-chain oxygen (Asp OD1, OD2; Glu OE1, OE2) on opposing chains fell within 4.0 Å. A hydrogen-bond-like contact required a donor–acceptor heavy-atom distance ≤ 3.5 Å together with a donor–hydrogen–acceptor angle ≥ 135°, evaluated in both donor directions. A hydrophobic contact required side-chain heavy atoms of two apolar residues (Ala, Val, Leu, Ile, Met, Phe, Trp, Tyr, Pro, Cys; backbone atoms excluded) within 4.5 Å. Aromatic and cation–π contacts used a 6.0 Å centroid cutoff.

The proximity term maps the mean minimum heavy-atom approach distance *d_i_*, averaged over contacting frames, linearly onto [0, 1] between 2.8 Å and the 4.5 Å contact cutoff,

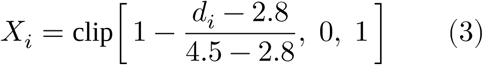

with *d_i_* set to the contact cutoff for residues that never contact Rac1.

The score therefore ranks residues by how consistently they touch Rac1. We call it an *interface-contact score*. We keep the term *hot spot* for its established energetic meaning: a residue whose substitution costs a defined amount of binding free energy (Bogan & Thorn, 1998). Scores were written to the PDB B-factor column for visualization in PyMOL (Schrödinger, LLC, 2025).

This identified an interface patch on residues 304–310 (Arg304, Val305, Ser306, Gly307, Phe308, Ser309, Asp310), which contains Arg304, the annotated catalytic arginine finger required for Rac GTPase activation (UniProt CHN1/P15882). Ser306 scored highest in the first trajectory, with Phe308 contributing aromatic surface. A second element comprised of Ile420, Val421, Pro424, Thr425, Ala434, Met435, Leu438, Ile441 and Arg445, with Ile420 ranking second in that trajectory. Electrostatic contacts Lys344, Asp310, Glu300, Arg348 and Glu334 were also retained with Lys344 observed to form a persistent salt bridge as Rac1 Asp63/Glu62 and Asp310 contacted Rac1 Arg94 (Supplementary Table S2; Fig. 2B,C).

**Figure 2.**
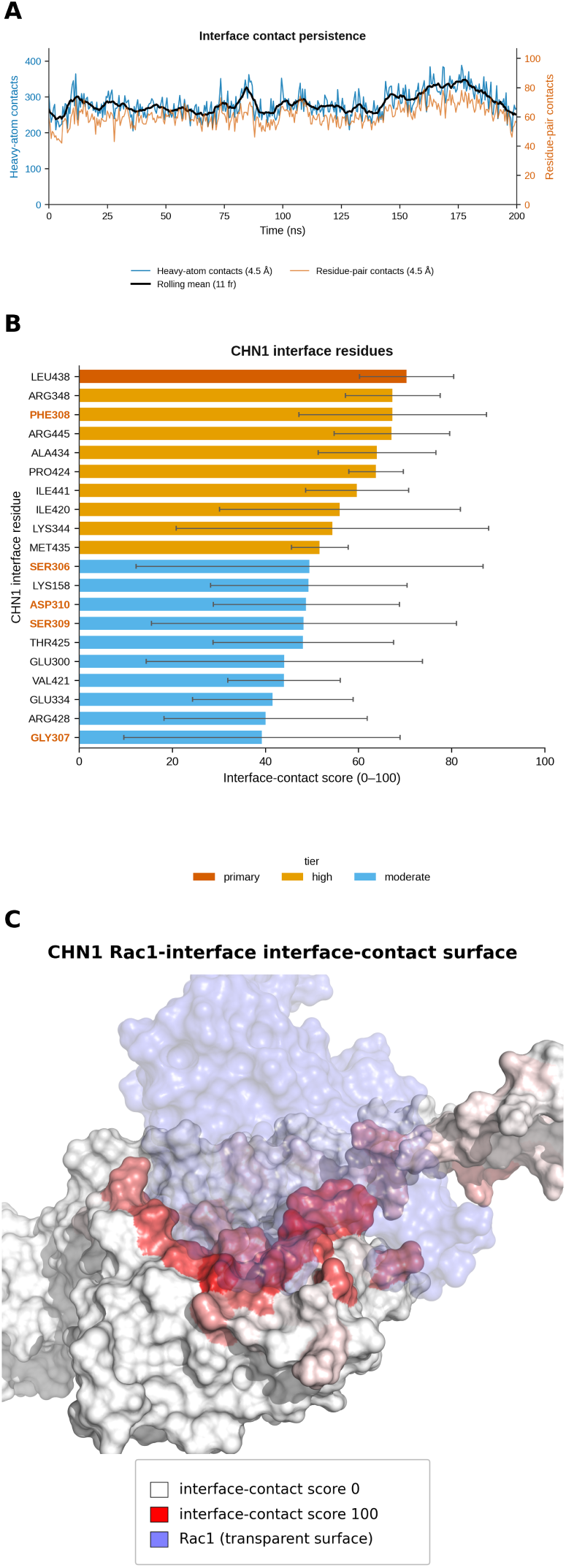
The CHN1–Rac1 interface resolves to a contiguous surface whose reproducible core is the Ile420–Arg445 ridge. **(A)** Interface contact persistence across the CHN1–Rac1 trajectory (replicate 1 shown). **(B)** Top-20 CHN1 interface residues ranked by the mean trajectory-derived interface-contact score over three independent 200 ns replicates; bars are the mean and error bars the inter-replicate SD (n = 3). Residues of the 304–310 patch are labeled in bold red: they rank below the Ile420–Arg445 ridge on the replicate mean and carry roughly twice its spread.

This residue set, centered on the Ile420–Arg445 ridge with the 304–310 patch and the neighboring electrostatic contacts, defined the CHN1-side interface used in subsequent work to characterize the interface dynamics.

### AlphaFold3 modeling of reported CHN1 interactors and ipSAE interface scoring

To contextualize the confidence of the CHN1–Rac1 model, we modeled CHN1 with its reported physical interactors from the BioGRID record for CHN1 (gene 107547). After removing duplicates, this yielded 31 unique interactors (Oughtred et al., 2021). Each CHN1–interactor pair was predicted using the AlphaFold3 server, which generated five models per complex (Abramson et al., 2024).

We calculated ipSAE for all five models using interchain residue pairs with PAE < 10 Å and Cβ–Cβ distance < 10 Å (Cα for glycine), then selected the model with the best ipSAE (d0dom) for each interactor. The selected models were evaluated using ipSAE, ipTM, pTM, chain-pair ipTM, and the number of confident interchain contacts (Dunbrack, 2025). Three high-confidence partners were clearly separated from the remaining interactors, with ipSAE values of approximately 0.45–0.68 compared with values near zero for the weak class (Supplementary Table S6).

### Molecular dynamics simulations and trajectory analysis

MD was applied at three stages: interface definition for the CHN1–Rac1 complex (above), and validation of the CHN1–interactor and of the CHN1–small-molecule and CHN1–peptide complexes. Crystallographic waters were removed, hydrogens added at pH 7.4, and each complex solvated under periodic boundary conditions in a TIP3P box (Jorgensen et al., 1983) with 1.0 nm minimum solute–wall padding, neutralizing counter ions and 0.15 M NaCl. CHN1–interactor and CHN1–peptide assemblies used CHARMM36 (July 2022 revision) (Huang & MacKerell, 2013); CHN1–small-molecule complexes used FF14SB (Maier et al., 2015) with ligands parameterized by GAFF2 and AM1-BCC charges (Jakalian et al., 2002; Wang et al., 2004) via ACPYPE (Sousa da Silva & Vranken, 2012), receptor and ligand topologies merged with ParmEd. Both structural CHN1 zinc ions were retained and modeled as bonded, tetrahedrally coordinated sites, each ligated by three cysteines and one histidine (His206/Cys236/Cys239/Cys255 and His244/Cys219/Cys222/Cys247). The coordinating cysteines were treated as thiolates and the histidines Nε-protonated, so that Nδ1 coordinates the zinc. Explicit Zn–ligand bonds and angle terms were added after topology generation.

Simulations were run with GROMACS 2025.3 (Abraham et al., 2015). Systems were energy-minimized by steepest descent, to a maximum force below 500 kJ mol^-1^ nm^-1^ within 50,000 steps. They were then given Maxwell–Boltzmann velocities at 310 K and equilibrated for 100 ps of NVT at 310 K, with solute heavy atoms restrained at 1000 kJ mol^-1^ nm^-2^ (≈2.4 kcal mol^-1^ Å^-2^). Small-molecule systems had a further 50 ps NVT warm-up at 1 fs. A 1000 ps NPT equilibration at 310 K and 1 atm retained the restraints and seeded unrestrained NPT production. A 4 fs time step was used, permitted by hydrogen-mass repartitioning (HMR) of the solute (Balusek et al., 2019), with frames every 500 ps, a 12 Å nonbonded cut-off, van der Waals force-switched between 10 and 12 Å, particle-mesh Ewald (PME) electrostatics and all bonds constrained by LINCS (Hess et al., 1997). The temperature was maintained with the velocity-rescaling thermostat (*τ*_T_ = 1.0 ps) and the pressure with the C-rescale (Eq2) and Parrinello–Rahman (Parrinello & Rahman, 1981) (Prod) barostats (*τ*_P_ = 5.0 ps).

The three top-ranked CHN1–interactor complexes (CHN1–Rac1, CHN1–NCK1, CHN1–NCK2), taken from the AlphaFold3 top models, were simulated in triplicate for 200 ns per replicate, each pairing the 445-residue CHN1 construct (UniProt P15882 residues 15–459, renumbered 1–445 throughout) with one full-length interactor (Rac1, 192 residues; NCK1, 377 residues; NCK2, 380 residues). Apo CHN1 was simulated identically in triplicate as the free-state reference. Sixteen CHN1–small-molecule complexes from the prioritized panel (ten generatively optimized analogues, five potency-optimized and five drug-likeness-optimized, plus six diverse natural-product leads) were each run for 100 ns across three compute nodes, and designed CHN1–peptide complexes for 100 ns in both macrocyclic and linear/linearized forms. For reproducibility, three independent 100 ns replicates were run for the prioritized CHN1–small-molecule complexes and for three peptide complexes, differing only in the random velocities assigned at the start of equilibration. Single-trajectory systems instead carry the per-frame standard deviation, labeled as such, since it measures sampling within one run. The production runs reported here total approximately 6.0 µs: triplicate 200 ns runs of the CHN1–Rac1, CHN1–NCK1, CHN1–NCK2 and apo-CHN1 systems (12 × 200 ns, the first CHN1–Rac1 replicate also serving as the interface-analysis trajectory); sixteen 100 ns CHN1–ligand complexes; eight 100 ns CHN1–peptide complexes; and the additional replicate runs of the three prioritized CHN1–ligand and three CHN1–peptide systems (12 × 100 ns), giving 6,000 ns in total. After periodic-boundary correction and alignment on the CHN1 backbone, RMSD, per-residue Cα RMSF, radius of gyration (Rg) and solvent-accessible surface area (SASA) were computed with GROMACS, and residue contacts with MDAnalysis (Gowers et al., 2016; Michaud-Agrawal et al., 2011). For the interactor complexes, target-fold stability was additionally quantified as the CHN1-core backbone RMSD (residues 1–445, fit and measured on the core), isolating the fold from the large-amplitude inter-domain motion of the multidomain NCK adaptors, with RMSF reported over both the whole trajectory and the last 10 ns. Interfaces were characterized by ligand/peptide center-of-mass displacement, contact persistence and interface-contact retention. Hydrogen bonds were detected per frame on a geometric criterion, giving protein–ligand counts for the lead complex and the CHN1 intramolecular count, alongside secondary structure assigned by DSSP (Kabsch & Sander, 1983) for the apo, Rac1-bound and lead-ligand-bound states, giving a free-versus-bound comparison of the fold (Supplementary Fig. S4; Table S6).

### Essential dynamics and free-energy landscapes

Collective backbone motion was extracted by principal component analysis of the Cα coordinates, diagonalizing the covariance matrix and projecting onto the first two eigenvectors with the GROMACS analysis tools. Fit and analysis used the same 445-atom CHN1 Cα selection in every system, so the analysis reports motion of the target rather than of the partner. Each trajectory was diagonalized separately, giving every state and every replicate its own eigenvector basis. Quantities compared between the free and bound states are restricted to basis-independent summaries of each covariance spectrum, namely the fraction of variance per mode, the mode count reaching 80 % of the variance, and the reproducibility of those quantities across replicates. The variance carried by each mode was taken from the eigenvalue spectrum, and the number of modes reaching 80 % of the variance recorded as a measure of how collective the motion is.

Gibbs free-energy landscapes over the first two components were built from the projected density at 310 K. The free energy of a bin follows from its probability P, normalized by the probability of the most populated bin:

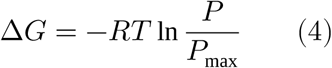

### End-point binding energies and per-residue decomposition

For each prioritized small-molecule complex, we calculated end-point binding energies and per-residue contributions from frames collected between 50 and 100 ns. For the peptide complexes we used frames from 25 to 100 ns. MM-PBSA calculations were performed with gmx_MMPBSA 1.6.4 across three 100 ns replicates and reported as the mean ± inter-replicate SD (Kollman et al., 2000; Miller et al., 2012; Valdés-Tresanco et al., 2021). The same trajectories were also analyzed by MM-GBSA using the OBC generalized Born model and 0.15 M salt. All calculations used the single-trajectory method at 310 K and 0.15 M ionic strength after correcting trajectories for periodic boundary conditions. The binding free energy (ΔG_binding_, kcal mol^-1^) was estimated from the following equation:

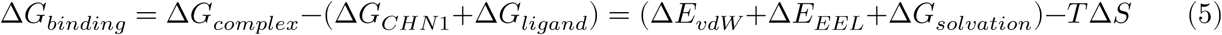

ΔE_vdW_ and ΔE_EEL_ are the molecular-mechanics van der Waals and electrostatic contributions, and ΔG_solvation_ the solvation free energy, partitioned into polar and nonpolar terms:

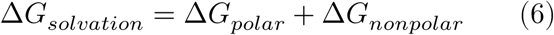

For MM-PBSA, the polar contribution was obtained by solving the Poisson–Boltzmann equation. The boundary between solute and solvent dielectric was defined from a level-set representation of the molecular surface. The solute and solvent dielectric constants were 1.0 and 80.0, the ionic strength 0.15 M, and the solvent probe radius 1.4 Å. The nonpolar contribution was modeled as a single term proportional to the solvent-accessible surface area.

Since Poisson–Boltzmann calculations were too costly for the larger protein–protein and peptide systems, their polar contributions were calculated with MM-GBSA using the OBC generalized Born model, mbondi2 radii, and 0.15 M salt (Onufriev et al., 2004). For CHN1–Rac1, each of the three replicate trajectories was analyzed from 50 to 200 ns. Interface free energies are reported as the mean ± inter-replicate SD.

Entropy changes (–TΔS) were estimated by interaction entropy (Duan et al., 2016) and C2 (Ekberg & Ryde, 2021) over the last 25 % of frames, normal-mode analysis being intractable at this size. The gas-phase interaction energy fluctuated by 7.1–9.7 kcal mol^-1^ for every ligand, above the ≈3.6 kcal mol^-1^ threshold for a reliable estimate. The reported ΔG_binding_ is therefore the enthalpic end-point term of Eq. (5), and the –TΔS term was likewise not evaluated for the protein–protein and protein–peptide systems. All values are consequently interpreted as relative end-point estimates (Genheden & Ryde, 2015).

Per-residue energy decomposition was calculated for all residues within 5 Å of the binding partner. This analysis identified the CHN1 residues that contributed most to binding, with negative values indicating favorable contributions and positive values indicating unfavorable contributions.

The two-dimensional interaction map for the top docking lead was generated with ProLIF and its LigNetwork module (Bouysset & Fiorucci, 2021; see Supplementary Figure S5). Because the refined protein structure lacked hydrogens, hydrogen bonds were assigned using heavy-atom donor and acceptor geometry.

## Results and discussion

### Structure prediction and molecular dynamics map candidate hot spots for the CHN1–Rac1 interface

We built the wild-type CHN1–Rac1 assembly by combining an AlphaFold3 prediction of the complex (pTM 0.70, ipTM 0.74) with the experimental CHN1 crystal structure (3CXL, 2.60 Å). Both structural zinc ions were retained (Fig. 1A; Methods) (Abramson et al., 2024; Shen et al., 2008). Relaxation by 200 ns of MD gave a stable complex whose interface could be defined dynamically. The contact map analysis was run on all three replicates and each score reported as a mean with inter-replicate SD (Fig. 2; Table S2).

The two elements behave differently across replicates. A hydrophobic ridge running from Ile420 to Arg445 is the reproducible core: Leu438 scores highest on the three-replicate mean (70.3 ± 10.1), with Arg445 (67.1 ± 12.4), Ala434 (64.0 ± 12.6), Pro424 (63.8 ± 5.9), Ile441 (59.7 ± 11.1), Ile420 (56.0 ± 25.9) and Met435 (51.7 ± 6.1), with Arg348 (67.4 ± 10.2) on the adjacent electrostatic edge. Its mean score is 59.2 with a mean inter-replicate SD of 12.6. The 304–310 patch scores lower and less consistently: Phe308 reaches 67.3 ± 20.1, but Ser306 averages 49.5 ± 37.2, Ser309 48.3 ± 32.7, Asp310 48.8 ± 20.0, and the catalytic arginine finger Arg304 falls outside the top twenty at 35.9 ± 26.3, rank 22 of 445. The patch mean is 45.1 with a mean inter-replicate SD of 28.0, against 12.6 for the ridge. One of the top-twenty residues, Lys158 (49.3 ± 21.1), lies inside the reconstructed 152–202 loop and is excluded from interpretation, its contacts being made by modeled residues.

Rank correlation between the ordering and the three-replicate mean was 0.85, and 18 of the top 20 residues were shared. Thus, the same interface region was consistently identified, although the exact residue ranking varied. In addition to these high-scoring contact regions, the interface contained a salt-bridge network, but its persistence varied between replicates. Lys344 formed a salt bridge with Rac1 Asp63/Glu62 that was almost continuous in two runs and nearly absent in the third, giving a mean frequency of 0.69 ± 0.51. Asp310 formed a salt bridge with Rac1 Arg94 in only one of the three runs, giving a mean frequency of 0.34 ± 0.48. Glu300, Arg348, and Glu334 made additional contacts of similarly mixed persistence (Supplementary Table S2). Mapping the interface-contact scores onto the CHN1 molecular surface revealed a shallow patch spanning the 304–310 loop and the Ile420 ridge (Fig. 2C). A Rac1 complex structure is not available, but the CHN1 side of the interface has been determined independently in both 3CXL and the 1.8 Å RhoGAP structure 2OSA. All 18 interface residues are resolved in both structures, which agree to 1.35 Å Cα RMSD across the domain and 1.00 Å over the high-scoring interface residues.

The AlphaFold3-reconstructed 152–202 loop lies outside the RhoGAP domain and is absent from 2OSA. This loop is likely intrinsically disordered and accounts for most of the RMSD and RMSF described below but is spatially separate from the interface.

The modeled CHN1-Rac1 orientation can be compared with other experimental results: no eukaryotic RhoGAP– Rac1 complex has been solved, but *Salmonella* SptP has been crystallized with Rac1 in a transition-state complex with GDP, Mg^2+^ and AlF_3_ (PDB 1G4U), fixing the catalytic geometry experimentally (Stebbins & Galán, 2000). Superposing our Rac1 on that of 1G4U (180 Cα, 0.94 Å RMSD) places Arg304 at the canonical arginine-finger locus, CZ 3.0 Å from SptP Arg209 CZ. This agreement is expected, because 1G4U and the other GAP–GTPase transition-state complexes predate the AlphaFold3 training cut-off. It also describes only the starting model. Arg304 begins 6.5 Å from the AlF_3_ center against 4.3 Å for SptP and withdraws to 10.9 ± 1.4 Å over the last 50 ns of the replicates as the vacant nucleotide site relaxes. The orientation itself remains modeled and a structure of the CHN1–Rac1 complex is the critical next step. The Rac1 switch regions remained close to active and transition-state conformations. Switch I and II were 1.1 and 0.6 Å from the active GMPPNP structure (1MH1) and 1.5 and 0.8 Å from the transition-state structure (1G4U), compared with 2.7 and 1.0 Å from the inactive GDP structure (1HH4). Because the active and transition-state structures differ by 1.6 Å at switch I, the modeled switches remained within the active conformational basin.

Arg304 illustrates the difference between catalytic importance and persistent interface binding. The contact score measures how consistently a residue engages the binding partner during the simulation. By contrast, an arginine finger enters the GTPase active site mainly during the transition state (Ahmadian et al., 1997; Kötting et al., 2008; Rittinger et al., 1997). In p50RhoGAP, replacing this residue reduces *k*_cat_ by more than 200-fold while having little effect on binding affinity (Graham et al., 1999). Arg304 is therefore not expected to behave as a conventional binding hotspot. Consistent with this, it scored highly in only one replicate, with scores of 66, 17, and 24 across the three simulations. Arg304 is best interpreted as a catalytic landmark, while the hydrophobic ridge forms the more persistent engagement surface.

None of the nine mapped CHN1 variants associated with DRS, Leu20Phe, Ile126Met, Tyr143His, Pro141Leu, Ala223Val, Gly228Ser, Pro252Gln, Pro252Ser, and Glu313Lys, came within 4 Å of Rac1 in any modeled complex (Fig. 1B; Supplementary Table S1). This supports the view that these mutations act allosterically by disrupting CHN1 autoinhibition (Colón-González et al., 2008; Miyake et al., 2008). The wild-type interface was therefore used to identify potential experimental probes.

Finally, we performed AlphaFold3 predictions of 31 BioGRID-reported CHN1 interactors with CHN1 itself, and scored each complex with an ipSAE interface-confidence metric (Supplementary Fig. S2; Supplementary Table S6). When the complexes were ranked by the domain-normalized ipSAE variant (ipSAE d0dom), only three rose above the weak class: NCK2 (0.79; rank 1, class Strong), NCK1 (0.78; rank 2, class Strong), and Rac1 (0.61; rank 3, class Good, and the highest AlphaFold3 ipTM of the whole set at 0.76), while the remaining 28 scored as weak (ipSAE ≈ 0). The recovery of Rac1 among the top three confident partners, alongside the two NCK adaptors that also bind through small folded interfaces, is consistent with the biological relevance of the modeled CHN1–Rac1 complex.

### Triplicate 200 ns simulations show a stable CHN1 core and persistent interactor interfaces

To compare the stability of the highest-confidence predicted interactions, each of the three CHN1 partners identified by the AlphaFold3 screen, Rac1, NCK1, and NCK2, was simulated in triplicate for 200 ns. The CHN1 core remained folded and conformationally stable in all nine trajectories (Supplementary Fig. S3), and the interfaces were largely maintained. The CHN1–Rac1 and CHN1–NCK1 complexes stayed fully associated in every replicate, whereas the CHN1–NCK2 complex remained associated in two of three runs and partially dissociated in the third (Supplementary Fig. S3). The CHN1–Rac1 replicates preserved the contact network defined above (Fig. 2A), most consistently the Ile420–Arg445 ridge, indicating that the surface is a persistent feature of the complex. Applying the MM-GBSA end-point method to each of the three CHN1–Rac1 replicates gave a mean interface end-point binding energy of −20.4 ± 10.0 kcal/mol (Supplementary Fig. S6B and Supplementary Table S4).

To determine whether Rac1 binding changes the collective motions of CHN1, we used principal component analysis of CHN1 Cα motion across three replicates of both the free and bound states (Fig. 3). Across all replicates, apo CHN1 required 6.3 ± 1.5 modes to explain 80% of the variance, compared with 5.3 ± 3.2 modes for the bound complex. The first two components explained 58.8 ± 7.9% of the motion in apo CHN1 and 66.3 ± 15.0% in the complex. Because these values overlap, we found no clear evidence that Rac1 binding concentrates CHN1 motion.

**Figure 3.**
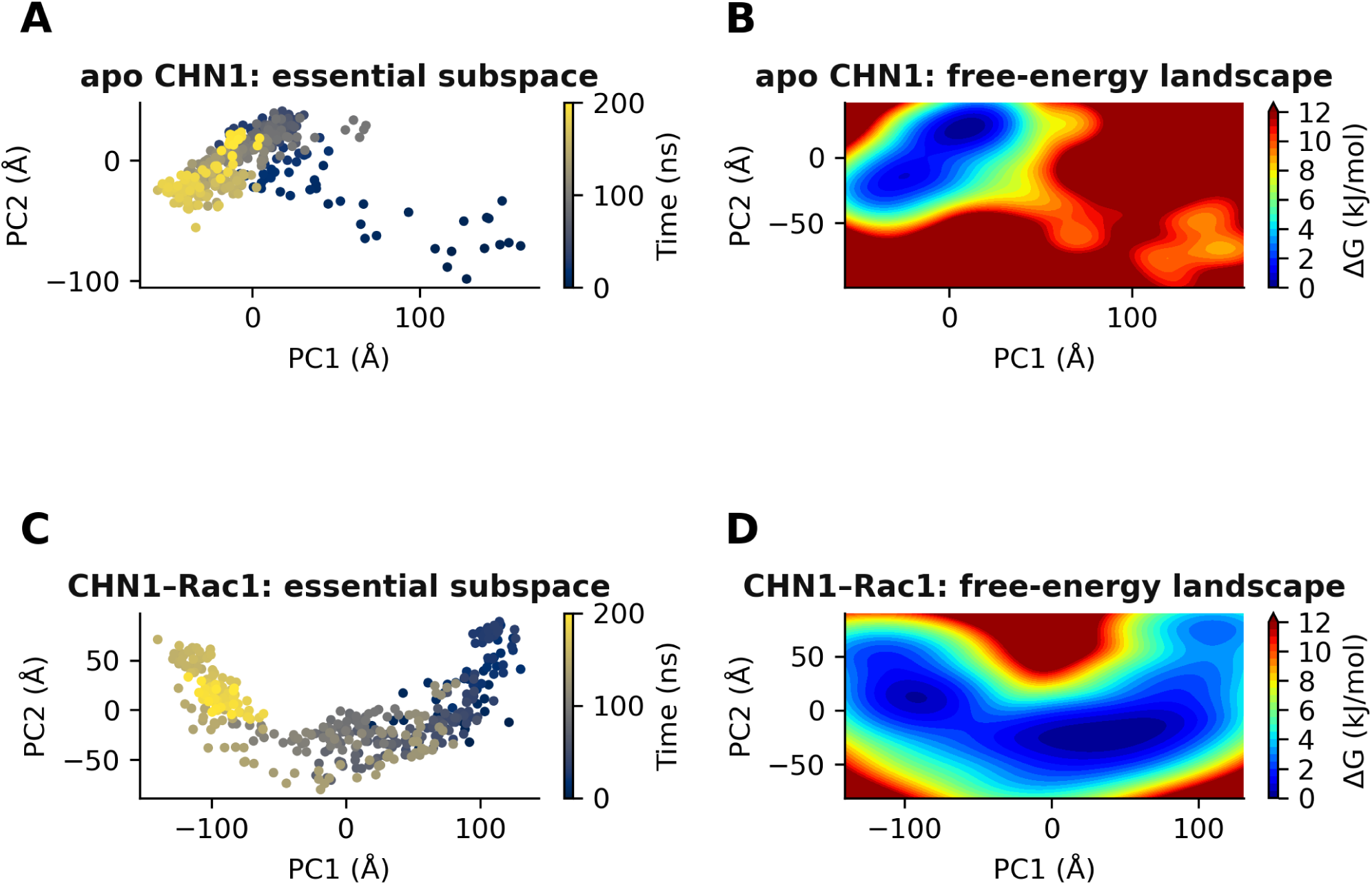
Rac1 binding does not reproducibly concentrate CHN1 backbone motion, but makes it markedly less reproducible. Essential dynamics of CHN1 free (apo) and within the CHN1–Rac1 complex, from principal component analysis of the CHN1 Cα coordinates computed identically for both states. Panels show replicate 1 of three for each. (A, B) Apo CHN1: projection onto the first two principal components colored by simulation time, and the corresponding Gibbs free-energy landscape (−RT ln P at 310 K). (C, D) The same for CHN1 within the complex. Across three replicates of each state the first two components carry 58.8 ± 7.9 % of the positional variance free and 66.3 ± 15.0 % bound, requiring 6.3 ± 1.5 and 5.3 ± 3.2 modes respectively to reach 80 %; these distributions overlap, so the concentration visible in this single pair of replicates is not reproducible.

Standard trajectory metrics showed that the CHN1–Rac1 complex remained compact, folded, and associated over 200 ns (Supplementary Fig. S4). The radius of gyration (30.0 ± 0.4 Å) and solvent-accessible surface area (342 ± 2 nm^2^) were stable across all three replicates, and Cα fluctuations remained low across the folded domains and high-scoring interface residues. The elevated backbone RMSD and dominant RMSF peak came entirely from the AlphaFold3-reconstructed 152–202 loop. Whole-backbone RMSD plateaued near 11 Å and averaged 8.7 ± 1.4 Å across the three runs (Table S7). Apo CHN1 showed a similar pattern, indicating that this motion is intrinsic to the modeled loop. Secondary structure and intramolecular hydrogen bonding were also similar in the free and bound states, showing that Rac1 binding did not disrupt the CHN1 fold (Supplementary Fig. S4; Table S6).

### Small molecules engage the dynamically defined CHN1–Rac1 interface

With Rac1 removed, the wild-type CHN1 interface was screened against the COCONUT natural-product library (Chandrasekhar et al., 2025) using Uni-Dock. Of 738,823 compounds, 547,719 were prepared and 250,371 were successfully docked. The top 5,000 were evaluated for chemical diversity, yielding 24 candidates. ADMET-AI filtering reduced these to eight Tier-1 leads and four rescue compounds (Fig. 4A). The Tier-1 leads bound near Ser306, Phe308, and the Lys344/Asp310 salt-bridge network, which are key parts of the Rac1 interface (Fig. 4B; Fig. S5). Their predicted affinities were moderate, with Vina scores of −6.5 to −7.1 kcal/mol and QED values of 0.67–0.91.

**Figure 4.**
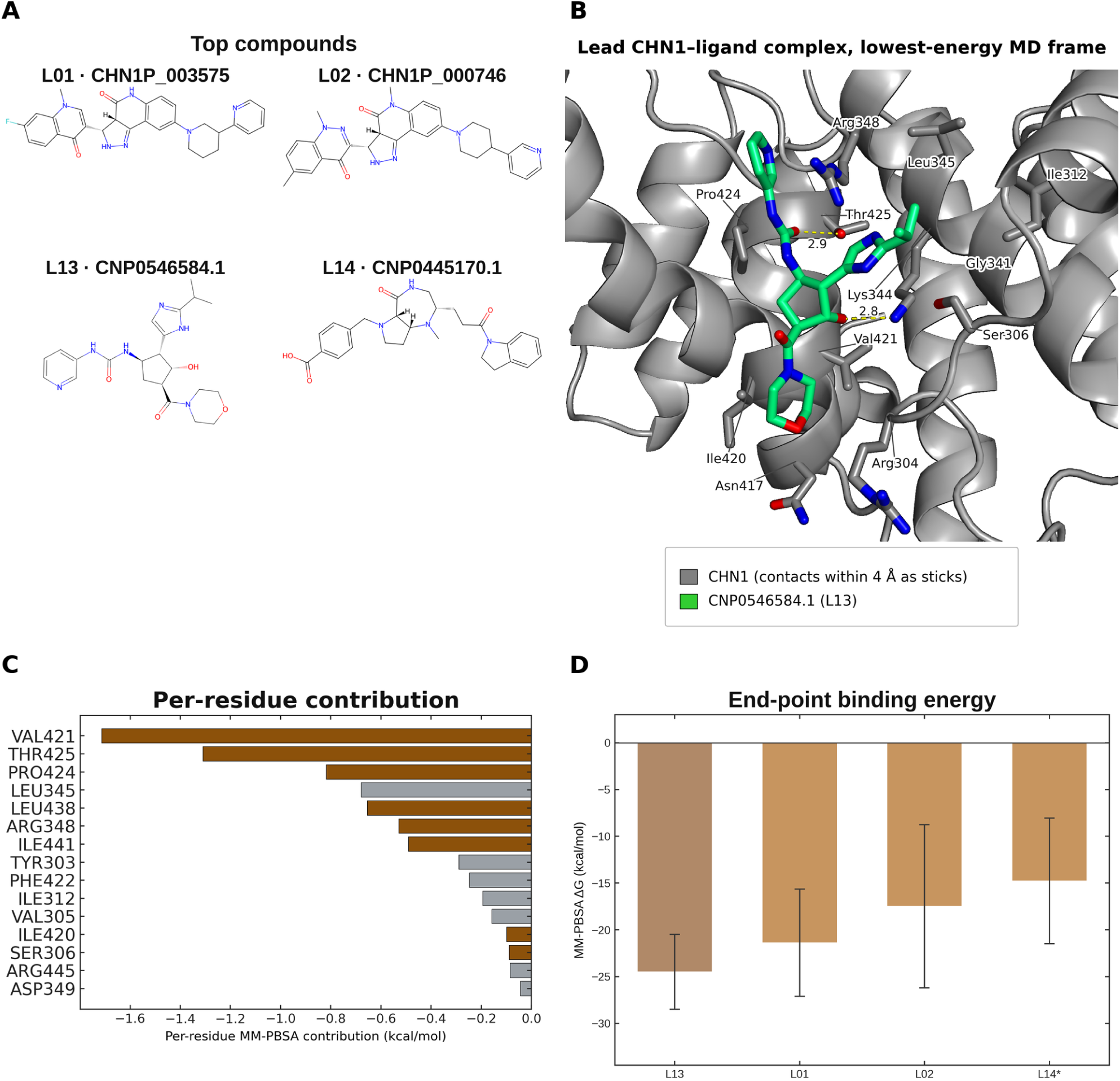
Small molecules placed on the interface-contact surface engage the residues that hold Rac1. **(A)** The four compounds carried into molecular dynamics (L01, L02, L13 and L14), labeled by compound ID; the full Tier-1 and CReM-optimized panel is given in Supplementary Fig. S8. **(B)** The prioritized lead CNP0546584.1 (L13) at the lowest-energy frame of its 100 ns trajectory (frame 145, 72 ns; MM-PBSA ΔG −36.0 kcal/mol); CHN1 residues within 4 Å are shown as sticks, with dashed lines marking polar contacts and their distances in Å. **(C)** Per-residue MM-PBSA energy decomposition for a representative CHN1–ligand complex, identifying the CHN1 high-scoring interface residues that contribute most to ligand binding. **(D)** MM-PBSA end-point binding energies for the CHN1–ligand complexes, with the lead CNP0546584.1 (L13) shown first (solid orange). Bars for L13, L01 and L02 are the mean over three independent 100 ns replicates with an inter-replicate standard deviation (n = 3); the L14 bar, marked with an asterisk, is a single trajectory with the per-frame standard deviation (n = 1). L13 gives the most favorable and most reproducible end-point score; L01 and L02 overlap within their standard deviations.

To improve binding to the shallow interface, the initial hits were expanded by CReM fragment growth using a larger 26 Å docking box. In total, 8,971 analogues were fast-docked, 336 were re-docked in triplicate, and ADMET filtering of the top 128 yielded 16 analogues with median Vina scores of ≤ −9.0 kcal/mol across three seeds (Fig. 4A). The strongest compound, CHN1P_003575 (L01), scored −9.8 kcal/mol, improving by 2.6 kcal/mol over its parent. CHN1P_000746 and CHN1P_003652 also scored −9.6 kcal/mol. Supplementary Table S8 links each L-number to its compound and SMILES. These optimized analogues had predicted DILI and hERG liabilities, so they were treated as pose probes. The natural products scored weaker but showed cleaner predicted ADMET profiles. The two approaches produced a chemically diverse panel of 16 compounds for MD analysis.

To test whether docked poses survived thermal fluctuation, prioritized CHN1–small-molecule complexes were advanced to MD and rescored by MM-PBSA. Production simulations of the panel showed that a subset of complexes maintained ligand occupancy at the interface-contact patch. The three prioritized complexes were each simulated in triplicate. This tested how reproducible the end-point estimates are. The binding energy (ΔG_binding_) from MM-PBSA is reported as a mean over the three replicates, with an inter-replicate standard deviation (Fig. 4D; Supplementary Table S3). The end-point energies were uniformly favorable (negative ΔG_binding_) and identified a consistent lead: the CNP0546584.1 complex was the most favorable and most reproducible end-point score at −24.5 ± 4.0 kcal/mol, ahead of two comparable analogues at −21.4 ± 5.7 and −17.5 ± 8.7 kcal/mol, with a weaker single-trajectory reference at −14.8 kcal/mol. The ranking of the lead is robust, since the same complex is strongest under both the MM-PBSA and the generalized-Born variant (MM-GBSA −34.5 ± 3.1 kcal/mol), whereas the two runners-up overlap within their standard deviations and are not reliably separable from one another. Additionally, individual single-trajectory values can be misleading: the strongest optimized analogue, CHN1P_003575 (L01), gave −27.9 kcal/mol from its first trajectory alone but fell to −21.4 ± 5.7 kcal/mol across replicates. More rigorous methods, including enhanced sampling and alchemical free-energy calculations, could provide more reliable estimates (Cournia et al., 2017; Wang et al., 2015). To understand the binding interface in more detail, we decomposed the end-point energies into the energetic contributions of individual residues at the interface-contact surface (Fig. 4C). This analysis attributed the dominant favorable ΔG_decomp_ contributions to the same CHN1 high-scoring interface residues identified from the CHN1–Rac1 trajectory, indicating that the prioritized ligands engage the intended interface.

### BoltzGen design of candidate peptide binders

As a complementary strategy better matched to a shallow protein–protein interface, we designed peptide binders directly against the CHN1 interface-contact surface (Fig. 5A). Constrained ∼12-residue cyclic macrocycles were generated with BoltzGen, conditioned on the 304–310 patch, in a large (≈10^4^-scale) interface-conditioned campaign (Passaro et al., 2025; Rettie et al., 2025; Stark et al., 2025). BoltzGen is a generative model that jointly designs backbones, assigns sequences by inverse folding, and folds and scores designs with an internal Boltz-2 model.

**Figure 5.**
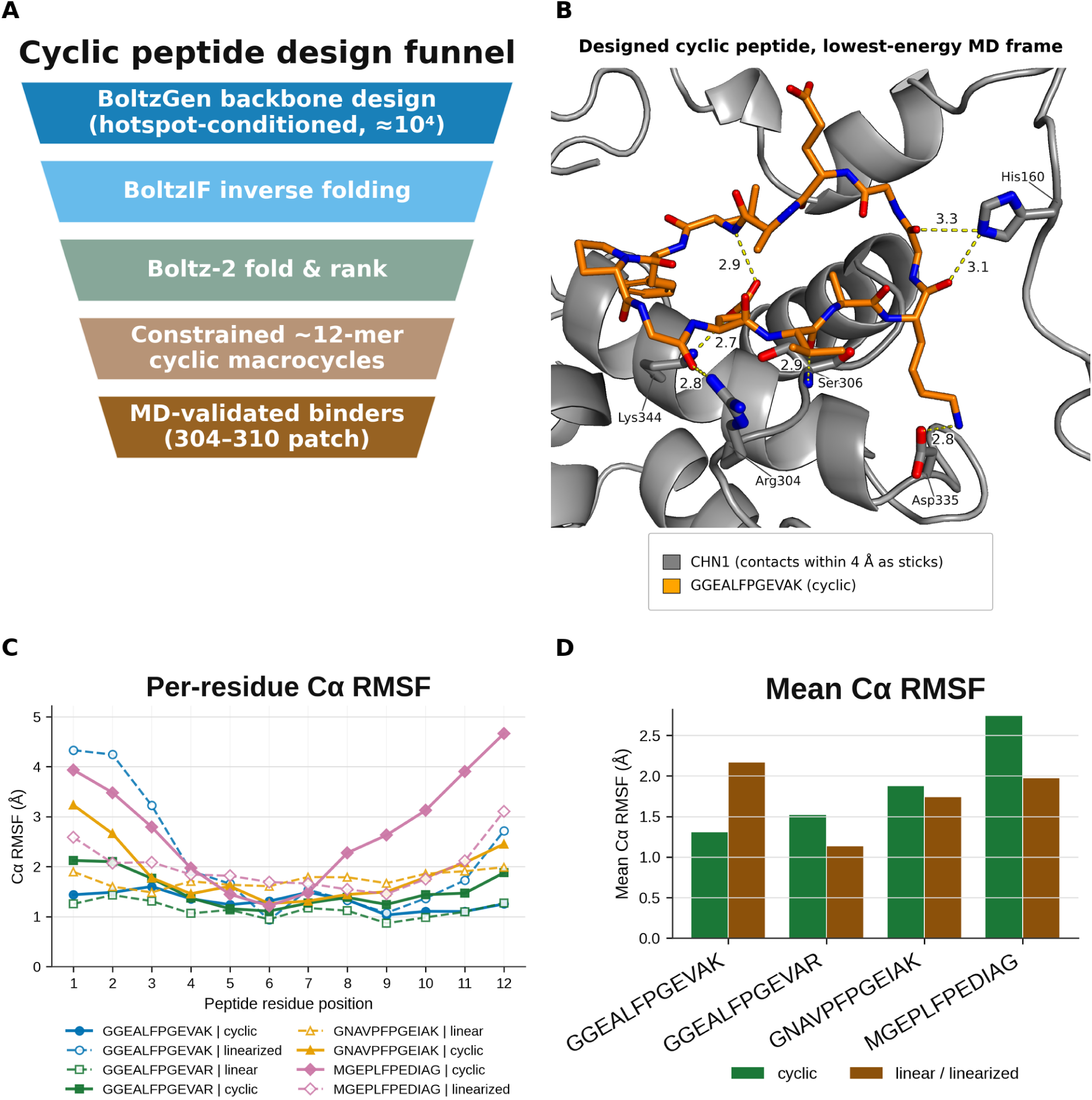
Diffusion-based cyclic-peptide design against the CHN1 interface-contact surface. **(A)** Peptide design funnel. **(B)** Designed cyclic peptide GGEALFPGEVAK at the lowest-energy frame of its 100 ns trajectory (frame 57, 28 ns; MM-GBSA ΔG −49.6 kcal/mol); CHN1 residues within 4 Å are shown as sticks, with dashed lines marking polar contacts and their distances in Å. **(C)** Mean last-10 ns Cα RMSF for the designed macrocyclic peptides.

MD of the designed peptides, performed for each sequence in both macrocyclic and linear/linearized forms, indicated that macrocyclization can substantially stabilize interface engagement, most clearly for matched cyclic/linear pairs (Fig. 5B,C). For the GGEALFPGEVAK design, the cyclic form maintained low per-residue Cα fluctuation over the last 10 ns of simulation (RMSF ≈ 1.0–1.6 Å across the peptide), whereas the corresponding linearized sequence fluctuated markedly more, particularly at the termini (RMSF up to ≈ 4.3 Å), and sampled the interface less persistently. Across the full panel, however, this cyclic-versus-linear advantage depended on the sequence. Some cyclic designs fluctuated as much as their linear counterparts, or more. One reason is that RMSF computed after alignment to the CHN1 receptor mixes internal peptide flexibility with rigid-body drift of the peptide away from the interface (Supplementary Table S5). Consistent with this interpretation, end-point MM-GBSA free energies computed for the matched cyclic and linear pairs were indistinguishable once replicate uncertainty was accounted for. For the GGEALFPGEVAR pair, each form was simulated in triplicate and rescored through an identical pipeline, giving −33.9 ± 6.1 kcal/mol for the macrocycle and −33.0 ± 6.9 kcal/mol for the linear counterpart (mean ± inter-replicate SD, n = 3); the difference of −0.85 kcal/mol is far smaller than the replicate spread. Comparing the two forms replicate-by-replicate (paired difference −0.85 ± 1.93 kcal/mol; cyclic−linear ≤ 3 kcal/mol in every replicate) indicates that the large per-design spread is trajectory-level noise shared by both constructs (Supplementary Fig. S6A and Supplementary Table S6). Macrocyclization therefore did not increase the raw end-point energy, and its benefit is expressed through improved interface retention and pre-organization (lower Cα RMSF). The constrained macrocyclic scaffolds are prioritized as the more promising interface-directed binders, particularly those that retained contacts with the CHN1 interface-contact surface (the 304–310 patch and/or the adjacent Ile420 ridge and the salt-bridge network) throughout the trajectories. This is consistent with the expectation that pre-organization reduces the entropic penalty of binding a flat surface (Vinogradov et al., 2019).

### Molecular dynamics indicates the interface can support stable small-molecule and peptide engagement

The prioritized lead small-molecule complex behaved comparably. In the first 100 ns trajectory of the best-scoring CHN1–ligand complex (CNP0546584.1; ΔG_MM-PBSA_ = −27.5 kcal/mol for this trajectory, −24.5 ± 4.0 kcal/mol across the three replicates), the protein core stayed folded, and the radius of gyration (27.6 ± 0.4 Å across the three replicates; Table S7) and SASA held constant. The ligand, meanwhile, relocated ∼11 Å from its initial docked pose over the first ∼15 ns, the docked pose being a weakly hydrogen-bonded, higher-energy starting point. It then settled into a distinct, MD-refined binding mode that it retained for the remainder of the trajectory (RMSD 1.3 ± 0.3 Å relative to the refined pose, in continuous pocket contact at a minimum ligand–pocket distance of ∼2 Å, engaged through three to four protein–ligand hydrogen bonds; Fig. 6B,D). The persistence of these interface hydrogen bonds and pocket contacts, along with the low fluctuation of the interface residues, supports the favorable endpoint free energy computed for this complex (Fig. 4D; Supplementary Table S3) and indicates stable engagement of the CHN1 interface-contact surface. That the ligand refined its pose substantially from the docking prediction is expected for a shallow protein–protein interface, where rigid docking is least reliable, and means the reported end-point free energies reflect the MD-relaxed geometry.

**Figure 6.**
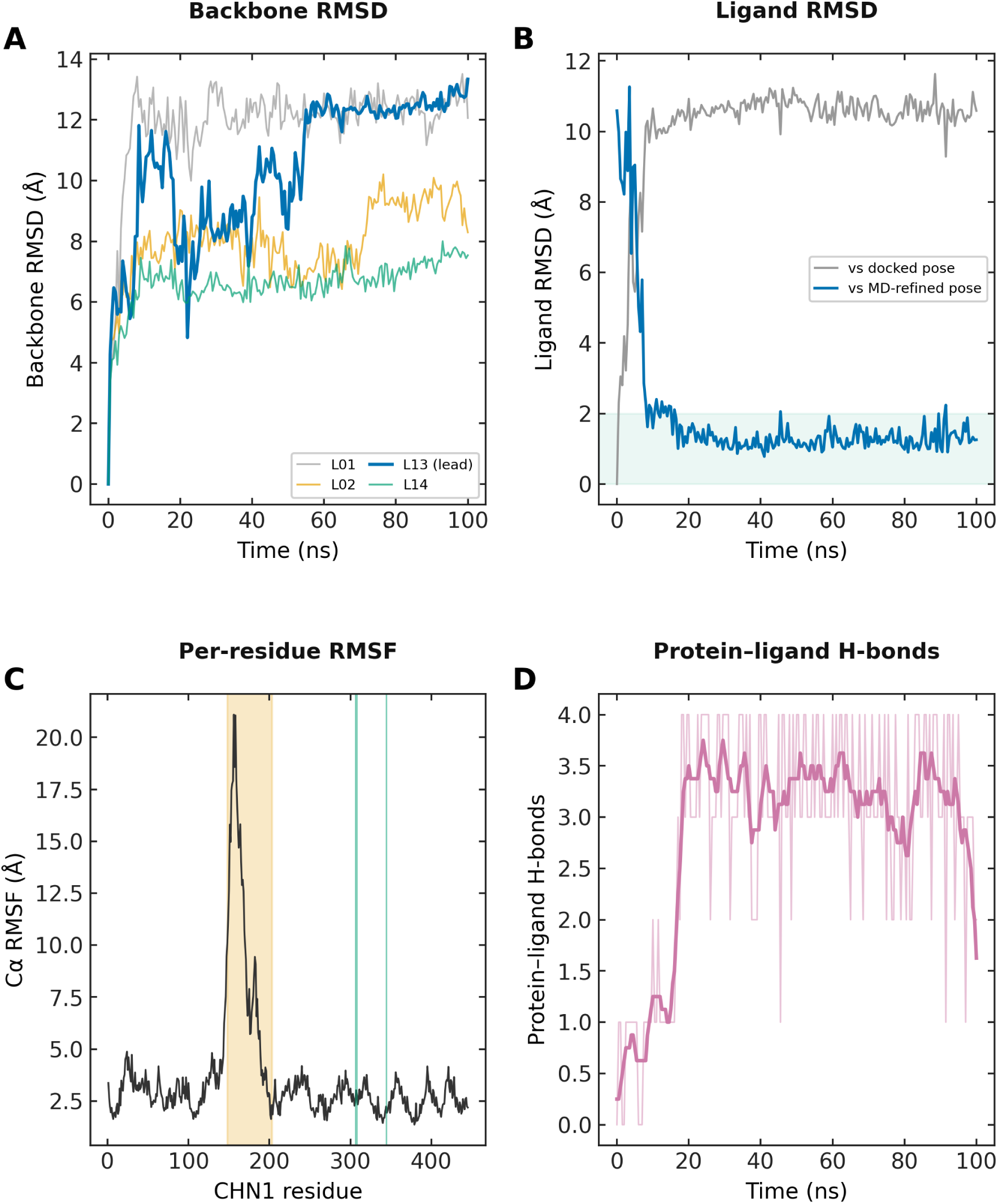
Molecular dynamics of the lead CHN1–ligand complex (CNP0546584.1; 100 ns). **(A)** Protein backbone RMSD versus time for the four MD-analyzed CHN1–ligand complexes (the lead CNP0546584.1 in bold). **(B)** Ligand heavy-atom RMSD (fit on the binding-pocket backbone) relative to the initial docked pose (grey) and to the MD-refined pose (blue, mean of 30–100 ns); the ligand relocates ∼11 Å from the docked pose within ∼15 ns and then settles into a stable refined pose (RMSD ≈ 1.3 Å; green band ≤ 2 Å), maintaining continuous pocket contact. **(C)** Protein per-residue Cα RMSF; fluctuation is confined to the AlphaFold3-rebuilt graft, residues 148–203 (shaded), while the high-scoring interface residues (marks at 306/308/344) remain rigid. **(D)** Number of protein–ligand hydrogen bonds versus time (thin line, per frame; thick line, running mean); the lead maintains three to four hydrogen bonds throughout, indicating persistent engagement of the CHN1 interface-contact surface.

Taken across ligand classes, the molecular-dynamics evidence indicates that the CHN1–Rac1 interface, though shallow, can support stable engagement by both small molecules and constrained peptides. Prioritized natural-product/optimized small molecules retained interface-contact occupancy and returned favorable end-point binding energies dominated by the same residues that dominate the Rac1 contact interface, and, for the one matched design pair simulated in both forms, the macrocycle was less flexible at the CHN1 interface-contact surface than its linearized counterpart (Cα RMSF 0.95 ± 0.02 versus 1.28 ± 0.09 Å). The convergence of two design modalities onto the same surface provides consistent support for the tractability of this interface as a ligand-binding surface.

### Interface tractability and the selectivity challenge

Gain-of-function CHN1 mutations hyperactivate α2-chimaerin, producing excessive RacGAP activity that disrupts ocular motor axon pathfinding (Miyake et al., 2008), but the interface through which this occurs had not been examined dynamically. Energetic mapping identified hot spots that could potentially be engaged (Wells & McClendon, 2007). However, energetic mapping alone cannot show whether these sites remain accessible on a flexible protein surface or whether a ligand can occupy them stably. This limitation motivated the small-molecule and peptide design studies.

The designed compounds remained bound during simulations and repeatedly contacted residues that also stabilize Rac1 binding. This suggests that the interface is not too shallow or mobile to be engaged (Rehman et al., 2023; Scott et al., 2016; Vassilev et al., 2004). These molecules provide potential experimental probes for testing whether the predicted interface residues can be occupied and perturbed.

The two molecular classes offer different experimental advantages. The small molecules had favorable predicted absorption and permeability profiles (Swanson et al., 2024) and may therefore be more useful for perturbing CHN1 in cellular systems. Macrocycles can engage broader and shallower surfaces that are difficult for conventional small molecules to capture (Dougherty et al., 2017) and may provide greater selectivity through their larger interaction footprint (Vinogradov et al., 2019). Because the axonal miswiring underlying DRS occurs early in development, reversing the disease process therapeutically would be difficult. These molecules could be used to test whether temporary disruption of the CHN1–Rac1 interaction changes Rac1 signalling or axon-guidance behavior.

Selectivity remains an important limitation. Rho-family signalling proteins share conserved surfaces, and the AlphaFold3 screen predicted CHN1 interactions with NCK1 and NCK2 at confidence levels comparable to or greater than Rac1. Any experimental probe would therefore need to be tested against these partners and related GAP–GTPase systems (Montalvo-Ortiz et al., 2012). Comparative modeling, mutational studies, and direct binding assays will be needed to determine whether the predicted compounds distinguish the Rac1 interface from other CHN1 interaction surfaces.

Rac1 dysregulation is involved in cancer, cardiovascular disease, and neurodegeneration, while chimaerins regulate cytoskeletal organization and cell signalling (Kazanietz & Caloca, 2017; Marei & Malliri, 2017; Yang & Kazanietz, 2007). Many of these pathways depend on transient and structurally flexible protein–protein interfaces that are difficult to characterize from static structures. The workflow used here combines structural modeling, replicate MD, energetic mapping, docking, and simulation-based ligand evaluation to identify persistent contacts and candidate probes for such interfaces.

These results define a persistent CHN1–Rac1 interaction surface and show that its central contact regions can be occupied by designed molecules. The next step is to test whether these compounds bind CHN1 experimentally and whether they perturb Rac1 engagement or signaling.

### Conclusions

By modeling a full length CHN1 construct using an experimental CHN1 crystal structure and an AlphaFold3 CHN1–Rac1 model and relaxing it by molecular dynamics, we identified the engagement surface with the Ile420– Arg445 hydrophobic ridge scoring highest and most consistently, while the 304–310 patch containing the catalytic arginine finger being conformationally variable. The disease mutations all lie more than 4 Å away, consistent with an allosteric mechanism. A complementary AlphaFold3/ipSAE interactor screen recovered Rac1 as a top-ranked CHN1 partner and simultaneously mapped the principal selectivity challenge (NCK1/NCK2). Principal component analysis across three replicates of each state showed no reproducible concentration of CHN1 backbone motion on binding (6.3 ± 1.5 modes to 80 % of the variance free against 5.3 ± 3.2 bound), correcting an apparent effect that a single pair of trajectories would have supported. Chemical sampling and cyclic-peptide design converged on binders occupying the same contact network, with end-point energies consistent with association being retained over the simulated interval. In the one matched pair simulated in triplicate, the macrocycle was less flexible than its linear form. While experimental validation remains essential, these results characterize the CHN1–Rac1 interface as a possible modulatory node and nominate a first set of candidate interface-directed molecular probes for interrogating it.

## Author contributions

**CRediT statement.** Conceptualization: Y.B. *Data curation:* A.S. *Formal analysis:* A.S. *Investigation:* A.S. *Methodology:* A.S., Y.B. *Project administration:* Y.B. *Resources:* Y.B. *Software:* A.S. *Supervision:* Y.B. *Validation:* A.S., Y.B. *Visualization:* A.S. *Writing, original draft:* A.S. *Writing, review and editing:* Y.B., A.S. Both authors read and approved the final version.

## Data availability statement

The original contributions presented in the study are included in the article and its Supplementary Material. Further inquiries can be directed to the corresponding author. Molecular-dynamics trajectories and the complete pipeline archive are available from the corresponding author on request.

## References

Abraham, M. J., Murtola, T., Schulz, R., Páll, S., Smith, J. C., Hess, B., & Lindahl, E. (2015). GROMACS: High performance molecular simulations through multi-level parallelism from laptops to supercomputers. SoftwareX, 1–2, 19–25. 10.1016/j.softx.2015.06.001

Abramson, J., Adler, J., Dunger, J., Evans, R., Green, T., Pritzel, A., Ronneberger, O., Willmore, L., Ballard, A. J., Bambrick, J., Bodenstein, S. W., Evans, D. A., Hung, C.-C., O’Neill, M., Reiman, D., Tunyasuvunakool, K., Wu, Z., Žemgulytė, A., Arvaniti, E., … Jumper, J. M. (2024). Accurate structure prediction of biomolecular interactions with AlphaFold 3. Nature, 630(8016), 493–500. 10.1038/s41586-024-07487-w

Ahmadian, M. R., Stege, P., Scheffzek, K., & Wittinghofer, A. (1997). Confirmation of the arginine-finger hypothesis for the GAP-stimulated GTP-hydrolysis reaction of Ras. Nature Structural Biology, 4(9), 686–689. 10.1038/nsb0997-686

Angelini, C., Trimouille, A., Arveiler, B., Espil-Taris, C., Ichinose, N., Lasseaux, E., Tourdias, T., & Lacombe, D. (2021). CHN1 and Duane retraction syndrome: Expanding the phenotype to cranial nerves development disease. European Journal of Medical Genetics, 64(4), Article 104188. 10.1016/j.ejmg.2021.104188

Aygit, E. D., Kocamaz, M., Inal, A., Fazil, K., Ocak, O. B., Akar, S., & Gokyigit, B. (2017). Management of Duane retraction syndrome with prismatic glasses. Clinical Ophthalmology, 11, 697–700. 10.2147/OPTH.S124183

Balusek, C., Hwang, H., Lau, C. H., Lundquist, K., Hazel, A., Pavlova, A., Lynch, D. L., Reggio, P. H., Wang, Y., & Gumbart, J. C. (2019). Accelerating membrane simulations with hydrogen mass repartitioning. Journal of Chemical Theory and Computation, 15(8), 4673–4686. 10.1021/acs.jctc.9b00160

Biler, E. D., Ilim, O., Onay, H., & Uretmen, O. (2017). CHN1 gene mutation analysis in patients with Duane retraction syndrome. Journal of AAPOS, 21(6), 472–475.e2. 10.1016/j.jaapos.2017.07.208

Bogan, A. A., & Thorn, K. S. (1998). Anatomy of hot spots in protein interfaces. Journal of Molecular Biology, 280(1), 1–9. 10.1006/jmbi.1998.1843

Bouysset, C., & Fiorucci, S. (2021). ProLIF: A library to encode molecular interactions as fingerprints. Journal of Cheminformatics, 13(1), 72. 10.1186/s13321-021-00548-6

Chan, W.-M., Miyake, N., Zhu-Tam, L., Andrews, C., & Engle, E. C. (2011). Two novel CHN1 mutations in 2 families with Duane retraction syndrome. Archives of Ophthalmology, 129(5), 649–652. 10.1001/archophthalmol.2011.84

Chandrasekhar, V., Rajan, K., Kanakam, S. R. S., Sharma, N., Weißenborn, V., Schaub, J., & Steinbeck, C. (2025). COCONUT 2.0: A comprehensive overhaul and curation of the collection of open natural products database. Nucleic Acids Research, 53(D1), D634–D643. 10.1093/nar/gkae1063

Clackson, T., & Wells, J. A. (1995). A hot spot of binding energy in a hormone-receptor interface. Science, 267(5196), 383–386. 10.1126/science.7529940

Colón-González, F., Leskow, F. C., & Kazanietz, M. G. (2008). Identification of an autoinhibitory mechanism that restricts C1 domain-mediated activation of the Rac-GAP α2-chimaerin. Journal of Biological Chemistry, 283(50), 35247–35257. 10.1074/jbc.M806264200

Cournia, Z., Allen, B., & Sherman, W. (2017). Relative binding free energy calculations in drug discovery: Recent advances and practical considerations. Journal of Chemical Information and Modeling, 57(12), 2911–2937. 10.1021/acs.jcim.7b00564

Cremer, J., Le, T., Ghahremanpour, M. M., Sługocka, E., Menezes, F., & Clevert, D.-A. (2026). FLOWR.ROOT – A flow matching-based foundation model for joint multi-purpose structure-aware 3D ligand generation and affinity prediction. Nature Communications, 17(1), Article 5883. 10.1038/s41467-026-74130-9

Dougherty, P. G., Qian, Z., & Pei, D. (2017). Macrocycles as protein–protein interaction inhibitors. Biochemical Journal, 474(7), 1109–1125. 10.1042/BCJ20160619

Duan, L., Liu, X., & Zhang, J. Z. H. (2016). Interaction entropy: A new paradigm for highly efficient and reliable computation of protein–ligand binding free energy. Journal of the American Chemical Society, 138(17), 5722–5728. 10.1021/jacs.6b02682

Dunbrack, R. L., Jr. (2025). Rēs ipSAE loquuntur: What’s wrong with AlphaFold’s ipTM score and how to fix it. bioRxiv. 10.1101/2025.02.10.637595

Eberhardt, J., Santos-Martins, D., Tillack, A. F., & Forli, S. (2021). AutoDock Vina 1.2.0: New docking methods, expanded force field, and Python bindings. Journal of Chemical Information and Modeling, 61(8), 3891–3898. 10.1021/acs.jcim.1c00203

Ekberg, V., & Ryde, U. (2021). On the use of interaction entropy and related methods to estimate binding entropies. Journal of Chemical Theory and Computation, 17(8), 5379–5391. 10.1021/acs.jctc.1c00374

Ferrario, J. E., Baskaran, P., Clark, C., Hendry, A., Lerner, O., Hintze, M., Allen, J., Chilton, J. K., & Guthrie, S. (2012). Axon guidance in the developing ocular motor system and Duane retraction syndrome depends on semaphorin signaling via α2-chimaerin. Proceedings of the National Academy of Sciences, 109(36), 14669–14674. 10.1073/pnas.1116481109

Genheden, S., & Ryde, U. (2015). The MM/PBSA and MM/GBSA methods to estimate ligand-binding affinities. Expert Opinion on Drug Discovery, 10(5), 449–461. 10.1517/17460441.2015.1032936

Gowers, R. J., Linke, M., Barnoud, J., Reddy, T. J. E., Melo, M. N., Seyler, S. L., Domański, J., Dotson, D. L., Buchoux, S., Kenney, I. M., & Beckstein, O. (2016). MDAnalysis: A Python package for the rapid analysis of molecular dynamics simulations. Proceedings of the 15th Python in Science Conference, 98–105. 10.25080/Majora-629e541a-00e

Graham, D. L., Eccleston, J. F., & Lowe, P. N. (1999). The conserved arginine in Rho-GTPase-activating protein is essential for efficient catalysis but not for complex formation with Rho·GDP and aluminum fluoride. Biochemistry, 38(3), 985–991. 10.1021/bi9821770

Hall, A., & Lalli, G. (2010). Rho and Ras GTPases in axon growth, guidance, and branching. Cold Spring Harbor Perspectives in Biology, 2(2), Article a001818. 10.1101/cshperspect.a001818

Hess, B., Bekker, H., Berendsen, H. J. C., & Fraaije, J. G. E. M. (1997). LINCS: A linear constraint solver for molecular simulations. Journal of Computational Chemistry, 18(12), 1463–1472. 10.1002/(SICI)1096-987X(199709)18:12<1463::AID-JCC4>3.0.CO;2-H

Huang, J., & MacKerell, A. D., Jr. (2013). CHARMM36 all-atom additive protein force field: Validation based on comparison to NMR data. Journal of Computational Chemistry, 34(25), 2135–2145. 10.1002/jcc.23354

Huber, A. (1974). Electrophysiology of the retraction syndromes. British Journal of Ophthalmology, 58(3), 293–300. 10.1136/bjo.58.3.293

Iwasato, T., Katoh, H., Nishimaru, H., Ishikawa, Y., Inoue, H., Saito, Y. M., Ando, R., Iwama, M., Takahashi, R., Negishi, M., & Itohara, S. (2007). Rac-GAP α-chimerin regulates motor-circuit formation as a key mediator of EphrinB3/EphA4 forward signaling. Cell, 130(4), 742–753. 10.1016/j.cell.2007.07.022

Jakalian, A., Jack, D. B., & Bayly, C. I. (2002). Fast, efficient generation of high-quality atomic charges. AM1BCC model: II. Parameterization and validation. Journal of Computational Chemistry, 23(16), 1623–1641. 10.1002/jcc.10128

Jorgensen, W. L., Chandrasekhar, J., Madura, J. D., Impey, R. W., & Klein, M. L. (1983). Comparison of simple potential functions for simulating liquid water. The Journal of Chemical Physics, 79(2), 926–935. 10.1063/1.445869

Jurgens, J. A., Barry, B. J., Chan, W.-M., MacKinnon, S., Whitman, M. C., Matos Ruiz, P. M., Pratt, B. M., England, E. M., Pais, L., Lemire, G., Groopman, E., Glaze, C., Russell, K. A., Singer-Berk, M., Di Gioia, S. A., Lee, A. S., Andrews, C., Shaaban, S., Wirth, M. M., … Engle, E. C. (2025). Expanding the genetics and phenotypes of ocular congenital cranial dysinnervation disorders. Genetics in Medicine, 27(4), 101216. 10.1016/j.gim.2024.101216

Kabsch, W., & Sander, C. (1983). Dictionary of protein secondary structure: Pattern recognition of hydrogen-bonded and geometrical features. Biopolymers, 22(12), 2577–2637. 10.1002/bip.360221211

Kazanietz, M. G., & Caloca, M. J. (2017). The Rac GTPase in cancer: From old concepts to new paradigms. Cancer Research, 77(20), 5445–5451. 10.1158/0008-5472.CAN-17-1456

Kekunnaya, R., & Negalur, M. (2017). Duane retraction syndrome: Causes, effects and management strategies. Clinical Ophthalmology, 11, 1917–1930. 10.2147/OPTH.S127481

Khademi Dehkordi, M., Hoveida, L., & Fani, N. (2024). Structure-based virtual screening, molecular docking, and molecular dynamics simulation approaches for identification of new potential inhibitors of class A β-lactamase enzymes. Journal of Biomolecular Structure and Dynamics, 42(11), 5631–5641. 10.1080/07391102.2023.2227724

Kollman, P. A., Massova, I., Reyes, C., Kuhn, B., Huo, S., Chong, L., Lee, M., Lee, T., Duan, Y., Wang, W., Donini, O., Cieplak, P., Srinivasan, J., Case, D. A., & Cheatham, T. E. (2000). Calculating structures and free energies of complex molecules: Combining molecular mechanics and continuum models. Accounts of Chemical Research, 33(12), 889–897. 10.1021/ar000033j

Kötting, C., Kallenbach, A., Suveyzdis, Y., Wittinghofer, A., & Gerwert, K. (2008). The GAP arginine finger movement into the catalytic site of Ras increases the activation entropy. Proceedings of the National Academy of Sciences, 105(17), 6260–6265. 10.1073/pnas.0712095105

Maier, J. A., Martinez, C., Kasavajhala, K., Wickstrom, L., Hauser, K. E., & Simmerling, C. (2015). ff14SB: Improving the accuracy of protein side chain and backbone parameters from ff99SB. Journal of Chemical Theory and Computation, 11(8), 3696–3713. 10.1021/acs.jctc.5b00255

Marei, H., & Malliri, A. (2017). Rac1 in human diseases: The therapeutic potential of targeting Rac1 signaling regulatory mechanisms. Small GTPases, 8(3), 139–163. 10.1080/21541248.2016.1211398

Michaud-Agrawal, N., Denning, E. J., Woolf, T. B., & Beckstein, O. (2011). MDAnalysis: A toolkit for the analysis of molecular dynamics simulations. Journal of Computational Chemistry, 32(10), 2319–2327. 10.1002/jcc.21787

Miller, B. R., III, McGee, T. D., Jr., Swails, J. M., Homeyer, N., Gohlke, H., & Roitberg, A. E. (2012). MMPBSA.py: An efficient program for end-state free energy calculations. Journal of Chemical Theory and Computation, 8(9), 3314–3321. 10.1021/ct300418h

Miyake, N., Chilton, J., Psatha, M., Cheng, L., Andrews, C., Chan, W.-M., Law, K., Crosier, M., Lindsay, S., Cheung, M., Allen, J., Gutowski, N. J., Ellard, S., Young, E., Iannaccone, A., Appukuttan, B., Stout, J. T., Christiansen, S., Ciccarelli, M. L., … Engle, E. C. (2008). Human CHN1 mutations hyperactivate α2-chimaerin and cause Duane’s retraction syndrome. Science, 321(5890), 839–843. 10.1126/science.1156121

Miyake, N., Demer, J. L., Shaaban, S., Andrews, C., Chan, W.-M., Christiansen, S. P., Hunter, D. G., & Engle, E. C. (2011). Expansion of the CHN1 strabismus phenotype. Investigative Ophthalmology & Visual Science, 52(9), 6321–6328. 10.1167/iovs.11-7950

Montalvo-Ortiz, B. L., Castillo-Pichardo, L., Hernández, E., Humphries-Bickley, T., De La Mota-Peynado, A., Cubano, L. A., Vlaar, C. P., & Dharmawardhane, S. (2012). Characterization of EHop-016, novel small molecule inhibitor of Rac GTPase. Journal of Biological Chemistry, 287(16), 13228–13238. 10.1074/jbc.M111.334524

Naik, B., Gupta, N., Godara, P., Srivastava, V., Kumar, P., Giri, R., Prajapati, V. K., Pandey, K. C., & Prusty, D. (2024). Structure-based virtual screening approach reveals natural multi-target compounds for the development of antimalarial drugs to combat drug resistance. Journal of Biomolecular Structure and Dynamics, 42(14), 7384–7408. 10.1080/07391102.2023.2240415

Newman, D. J., & Cragg, G. M. (2020). Natural products as sources of new drugs over the nearly four decades from 01/1981 to 09/2019. Journal of Natural Products, 83(3), 770–803. 10.1021/acs.jnatprod.9b01285

O’Boyle, N. M., Banck, M., James, C. A., Morley, C., Vandermeersch, T., & Hutchison, G. R. (2011). Open Babel: An open chemical toolbox. Journal of Cheminformatics, 3(1), Article 33. 10.1186/1758-2946-3-33

Onufriev, A., Bashford, D., & Case, D. A. (2004). Exploring protein native states and large-scale conformational changes with a modified generalized born model. Proteins: Structure, Function, and Bioinformatics, 55(2), 383–394. 10.1002/prot.20033

Oughtred, R., Rust, J., Chang, C., Breitkreutz, B.-J., Stark, C., Willems, A., Boucher, L., Leung, G., Kolas, N., Zhang, F., Dolma, S., Coulombe-Huntington, J., Chatr-aryamontri, A., Dolinski, K., & Tyers, M. (2021). The BioGRID database: A comprehensive biomedical resource of curated protein, genetic, and chemical interactions. Protein Science, 30(1), 187–200. 10.1002/pro.3978

Parrinello, M., & Rahman, A. (1981). Polymorphic transitions in single crystals: A new molecular dynamics method. Journal of Applied Physics, 52(12), 7182–7190. 10.1063/1.328693

Passaro, S., Corso, G., Wohlwend, J., Reveiz, M., Thaler, S., Somnath, V. R., Getz, N., Portnoi, T., Roy, J., Stark, H., Kwabi-Addo, D., Beaini, D., Jaakkola, T., & Barzilay, R. (2025). Boltz-2: Towards accurate and efficient binding affinity prediction. bioRxiv. 10.1101/2025.06.14.659707

Polishchuk, P. (2020). CReM: Chemically reasonable mutations framework for structure generation. Journal of Cheminformatics, 12(1), Article 28. 10.1186/s13321-020-00431-w

RDKit. (2025). RDKit: Open-source cheminformatics (Version 2025.09.1) [Computer software]. https://www.rdkit.org

Rehman, A. U., Khurshid, B., Ali, Y., Rasheed, S., Wadood, A., Ng, H. L., Chen, H. F., Wei, Z., Luo, R., & Zhang, J. (2023). Computational approaches for the design of modulators targeting protein–protein interactions. Expert Opinion on Drug Discovery, 18(3), 315–333. 10.1080/17460441.2023.2171396

Rettie, S. A., Juergens, D., Adebomi, V., Bueso, Y. F., Zhao, Q., Leveille, A. N., Liu, A., Bera, A. K., Wilms, J. A., Üffing, A., Kang, A., Brackenbrough, E., Lamb, M., Gerben, S. R., Murray, A., Levine, P. M., Schneider, M., Vasireddy, V., Ovchinnikov, S., … Bhardwaj, G. (2025). Accurate de novo design of high-affinity protein-binding macrocycles using deep learning. Nature Chemical Biology, 21(12), 1948– 1956. 10.1038/s41589-025-01929-w

Rittinger, K., Walker, P. A., Eccleston, J. F., Smerdon, S. J., & Gamblin, S. J. (1997). Structure at 1.65 Å of RhoA and its GTPase-activating protein in complex with a transition-state analogue. Nature, 389(6652), 758–762. 10.1038/39651

Schrödinger, LLC. (2025). The PyMOL molecular graphics system (Version 3.1.0) [Computer software]. https://www.pymol.org

Scott, D. E., Bayly, A. R., Abell, C., & Skidmore, J. (2016). Small molecules, big targets: Drug discovery faces the protein–protein interaction challenge. Nature Reviews Drug Discovery, 15(8), 533–550. 10.1038/nrd.2016.29

Shen, L., Buck, M., Tong, Y., Tempel, W., MacKenzie, F., Arrowsmith, C. H., Edwards, A. M., Bountra, C., Wilkstrom, M., Bochkarev, A., & Park, H. (2008). Crystal structure of human chimerin 1 (CHN1) [PDB entry 3CXL]. RCSB Protein Data Bank. 10.2210/pdb3CXL/pdb

Sheth, J., Ezisi, C. N., Tibrewal, S., Sachdeva, V., & Kekunnaya, R. (2021). Surgical outcomes of exotropic Duane retraction syndrome from a tertiary eye care center. Journal of Pediatric Ophthalmology and Strabismus, 58(1), 9–16. 10.3928/01913913-20200910-02

Sousa da Silva, A. W., & Vranken, W. F. (2012). ACPYPE AnteChamber PYthon Parser interfacE. BMC Research Notes, 5(1), 367. 10.1186/1756-0500-5-367

Stark, H., Faltings, F., Choi, M., Xie, Y., Hur, E., O’Donnell, T., Bushuiev, A., Uçar, T., Passaro, S., Mao, W., Reveiz, M., Bushuiev, R., Portnoi, T., Pluskal, T., Sivic, J., Kreis, K., Vahdat, A., Ray, S., Goldstein, J. T., … Jaakkola, T. (2025). BoltzGen: Toward universal binder design. bioRxiv. 10.1101/2025.11.20.689494

Stebbins, C. E., & Galán, J. E. (2000). Modulation of host signaling by a bacterial mimic: Structure of the *Salmonella* effector SptP bound to Rac1. Molecular Cell, 6(6), 1449–1460. 10.1016/S1097-2765(00)00141-6

Swanson, K., Walther, P., Leitz, J., Mukherjee, S., Wu, J. C., Shivnaraine, R. V., & Zou, J. (2024). ADMET-AI: A machine learning ADMET platform for evaluation of large-scale chemical libraries. Bioinformatics, 40(7), btae416. 10.1093/bioinformatics/btae416

Valdés-Tresanco, M. S., Valdés-Tresanco, M. E., Valiente, P. A., & Moreno, E. (2021). gmx_MMPBSA: A new tool to perform end-state free energy calculations with GROMACS. Journal of Chemical Theory and Computation, 17(10), 6281–6291. 10.1021/acs.jctc.1c00645

Vassilev, L. T., Vu, B. T., Graves, B., Carvajal, D., Podlaski, F., Filipovic, Z., Kong, N., Kammlott, U., Lukacs, C., Klein, C., Fotouhi, N., & Liu, E. A. (2004). In vivo activation of the p53 pathway by small-molecule antagonists of MDM2. Science, 303(5659), 844–848. 10.1126/science.1092472

Vinogradov, A. A., Yin, Y., & Suga, H. (2019). Macrocyclic peptides as drug candidates: Recent progress and remaining challenges. Journal of the American Chemical Society, 141(10), 4167–4181. 10.1021/jacs.8b13178

Walker, J. R., Hong, B. S., Shen, L., Arrowsmith, C. H., Sundstrom, M., Weigelt, J., Edwards, A. M., Bochkarev, A., & Park, H. W. (2007). *The Rho-GAP domain of human N-chimaerin* (PDB 2OSA) [Data set]. RCSB Protein Data Bank. 10.2210/pdb2OSA/pdb

Wang, J., Wolf, R. M., Caldwell, J. W., Kollman, P. A., & Case, D. A. (2004). Development and testing of a general amber force field. Journal of Computational Chemistry, 25(9), 1157–1174. 10.1002/jcc.20035

Wang, L., Wu, Y., Deng, Y., Kim, B., Pierce, L., Krilov, G., Lupyan, D., Robinson, S., Dahlgren, M. K., Greenwood, J., Romero, D. L., Masse, C., Knight, J. L., Steinbrecher, T., Beuming, T., Damm, W., Harder, E., Sherman, W., Brewer, M., … Abel, R. (2015). Accurate and reliable prediction of relative ligand binding potency in prospective drug discovery by way of a modern free-energy calculation protocol and force field. Journal of the American Chemical Society, 137(7), 2695–2703. 10.1021/ja512751q

Wells, J. A., & McClendon, C. L. (2007). Reaching for high-hanging fruit in drug discovery at protein–protein interfaces. Nature, 450(7172), 1001–1009. 10.1038/nature06526

Whitman, M. C., & Engle, E. C. (2017). Ocular congenital cranial dysinnervation disorders (CCDDs): Insights into axon growth and guidance. Human Molecular Genetics, 26(R1), R37–R44. 10.1093/hmg/ddx168

Yang, C., & Kazanietz, M. G. (2007). Chimaerins: GAPs that bridge diacylglycerol signalling and the small G-protein Rac. Biochemical Journal, 403(1), 1–12. 10.1042/BJ20061750

Yu, Y., Cai, C., Wang, J., Bo, Z., Zhu, Z., & Zheng, H. (2023). Uni-Dock: GPU-accelerated docking enables ultralarge virtual screening. Journal of Chemical Theory and Computation, 19(11), 3336–3345. 10.1021/acs.jctc.2c01145

Zhang, R., Jia, H., Chang, Q., Zhang, Z., Peng, C., Ma, Q., Liang, Y., Yang, S., & Jiao, Y. (2024). Two novel CHN1 variants identified in Duane retraction syndrome pedigrees disrupt development of ocular motor nerves in zebrafish. Journal of Human Genetics, 69(1), 33–39. 10.1038/s10038-023-01201-w

Zhou, T.-C., Duan, W.-H., Fu, X.-L., Zhu, Q., Guo, L.-Y., Zhou, Y., Hua, Z.-J., Li, X.-J., Yang, D.-M., Zhang, J.-Y., Yin, J., Zhang, X.-F., Zhou, G.-L., & Hu, M. (2020). Identification of a novel CHN1 p.(Phe213Val) variant in a large Han Chinese family with congenital Duane retraction syndrome. Scientific Reports, 10(1), 16225. 10.1038/s41598-020-73190-1

