## Supplementary Figures and Tables for "Dynamics of the α2-chimaerin–Rac1 interface in Duane retraction syndrome and computational design of candidate probes"

### Supplementary Information

#### Supplementary Figures

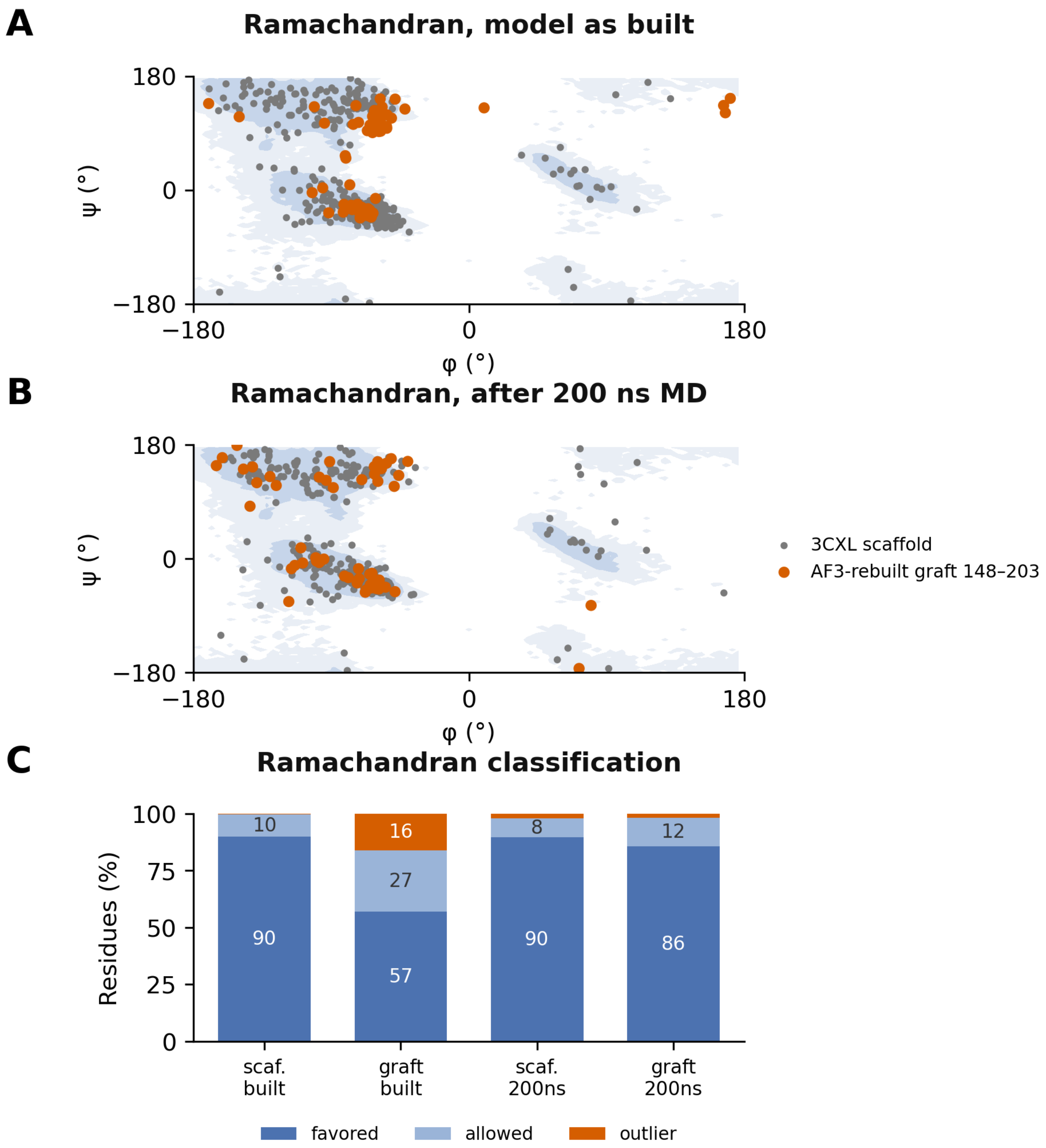


**Figure S1.** Construction and stereochemical validation of the hybrid CHN1 model. Ramachandran geometry of the 3CXL-derived scaffold (grey) and the AlphaFold3-rebuilt graft spanning residues 148–203 (red); shading is the MDAnalysis reference density at its favored and allowed contours. **(A)** As built. **(B)** After 200 ns of molecular dynamics. **(C)** Ramachandran classification of the four groups.

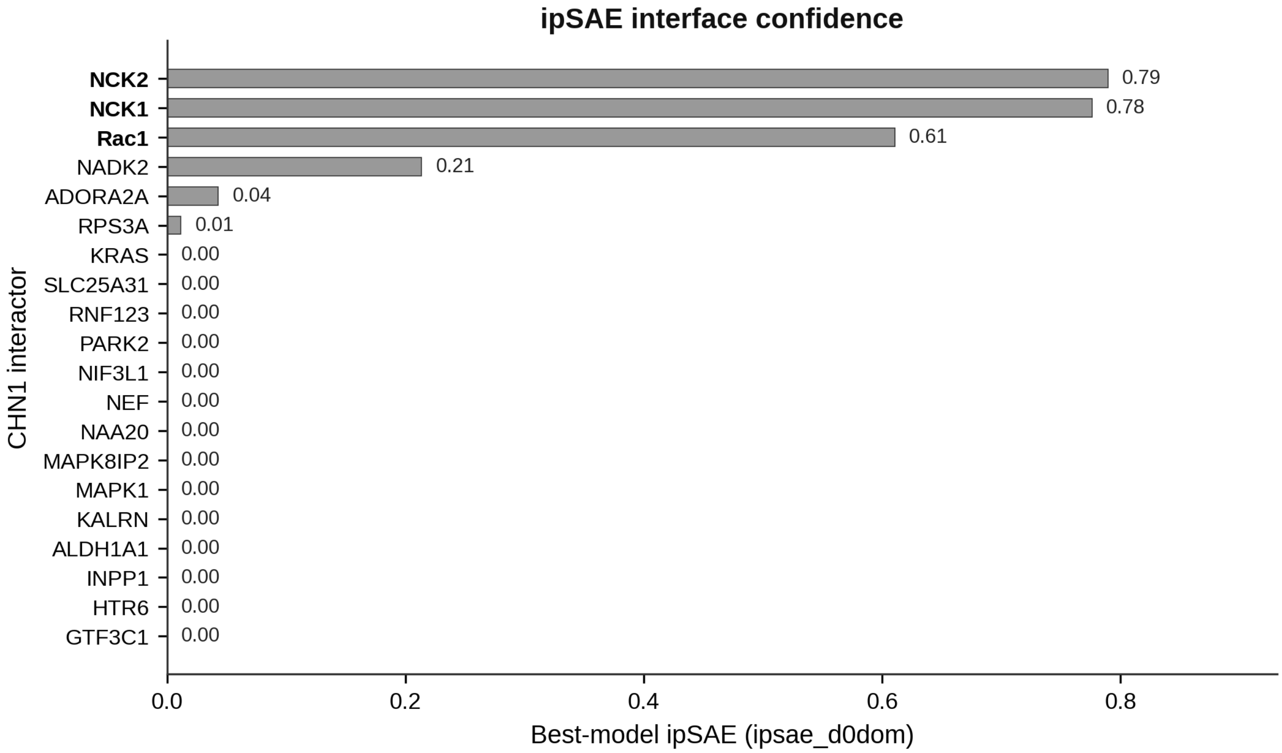


**Figure S2.** AlphaFold3 modeling of 31 BioGRID-reported CHN1 interactors, ranked by ipSAE interface confidence. NCK2, NCK1 and Rac1 are the only partners scoring above the confidence threshold.

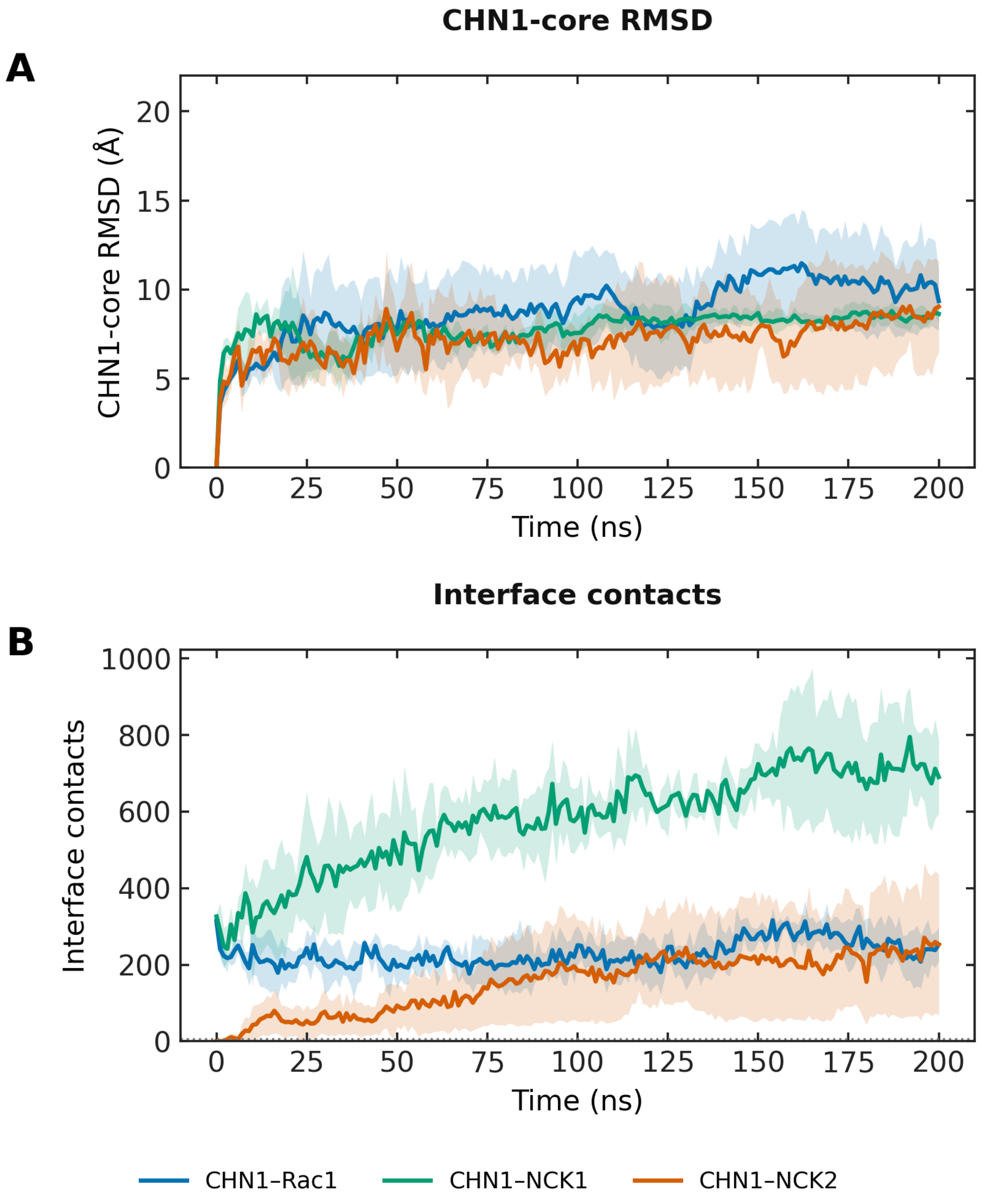


**Figure S3.** Stability of the three top-ranked CHN1-interactor complexes (CHN1–Rac1, CHN1–NCK1, CHN1–NCK2), as the mean ± SD over three independent 200 ns replicates (lines, mean; bands, ± SD). **(A)** CHN1-core backbone RMSD, fit and measured on the CHN1 core. **(B)** CHN1-core–interactor interface heavy-atom contacts (≤ 4.5 Å; dotted line).

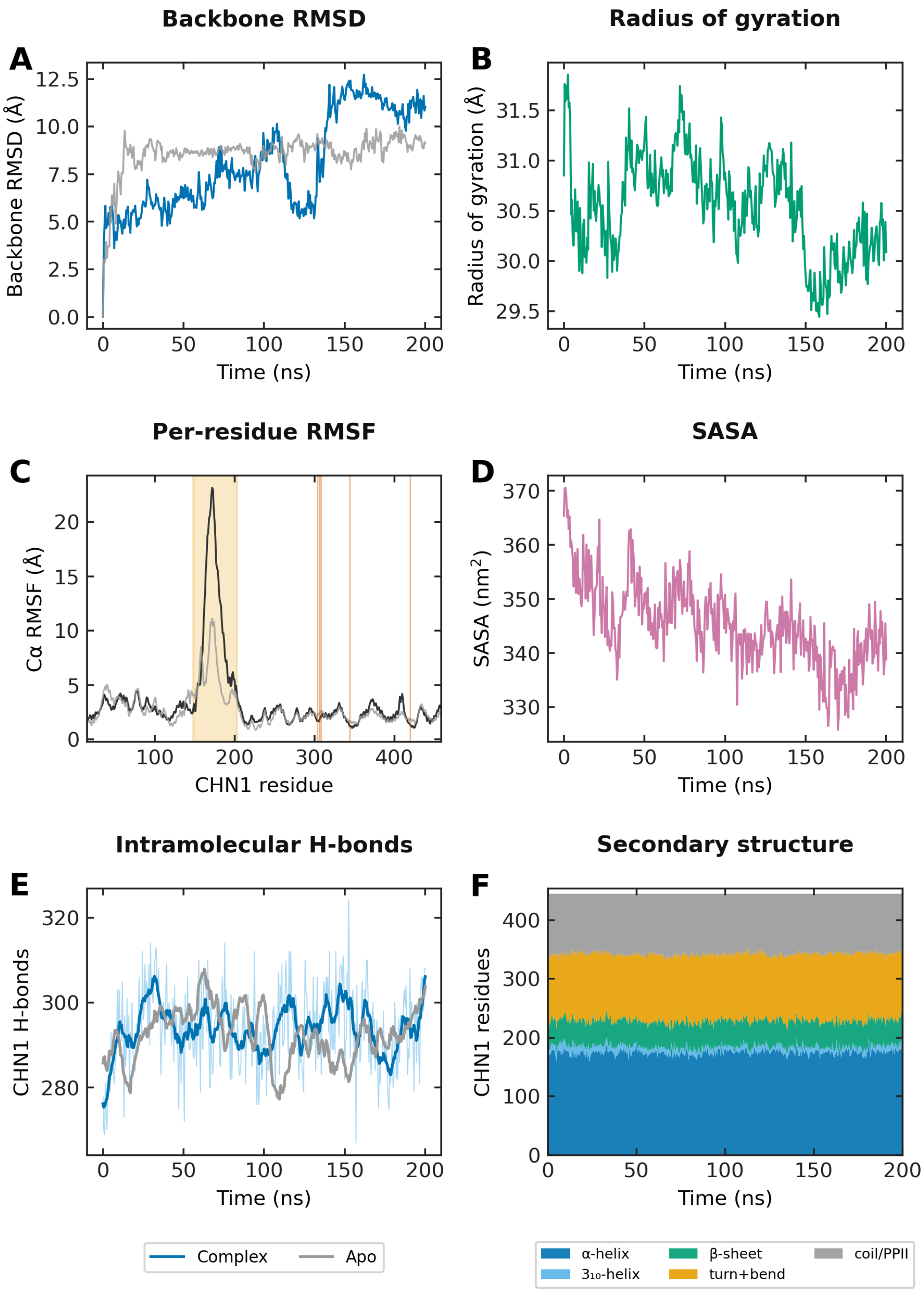


**Figure S4.** Molecular-dynamics metrics of the CHN1–Rac1 complex (replicate 1; 200 ns), with apo CHN1 overlaid where the comparison is like-for-like. **(A)** Backbone RMSD. **(B)** Radius of gyration. **(C)** Per-residue Cα RMSF for the complex (black) and apo CHN1 (grey); the AlphaFold3-rebuilt graft (148–203) is shaded and high-scoring interface residues are marked. **(D)** Solvent-accessible surface area. **(E)** CHN1 intramolecular hydrogen bonds. **(F)** DSSP secondary-structure content.

**Protein-ligand interaction map for CNP0062561.1**

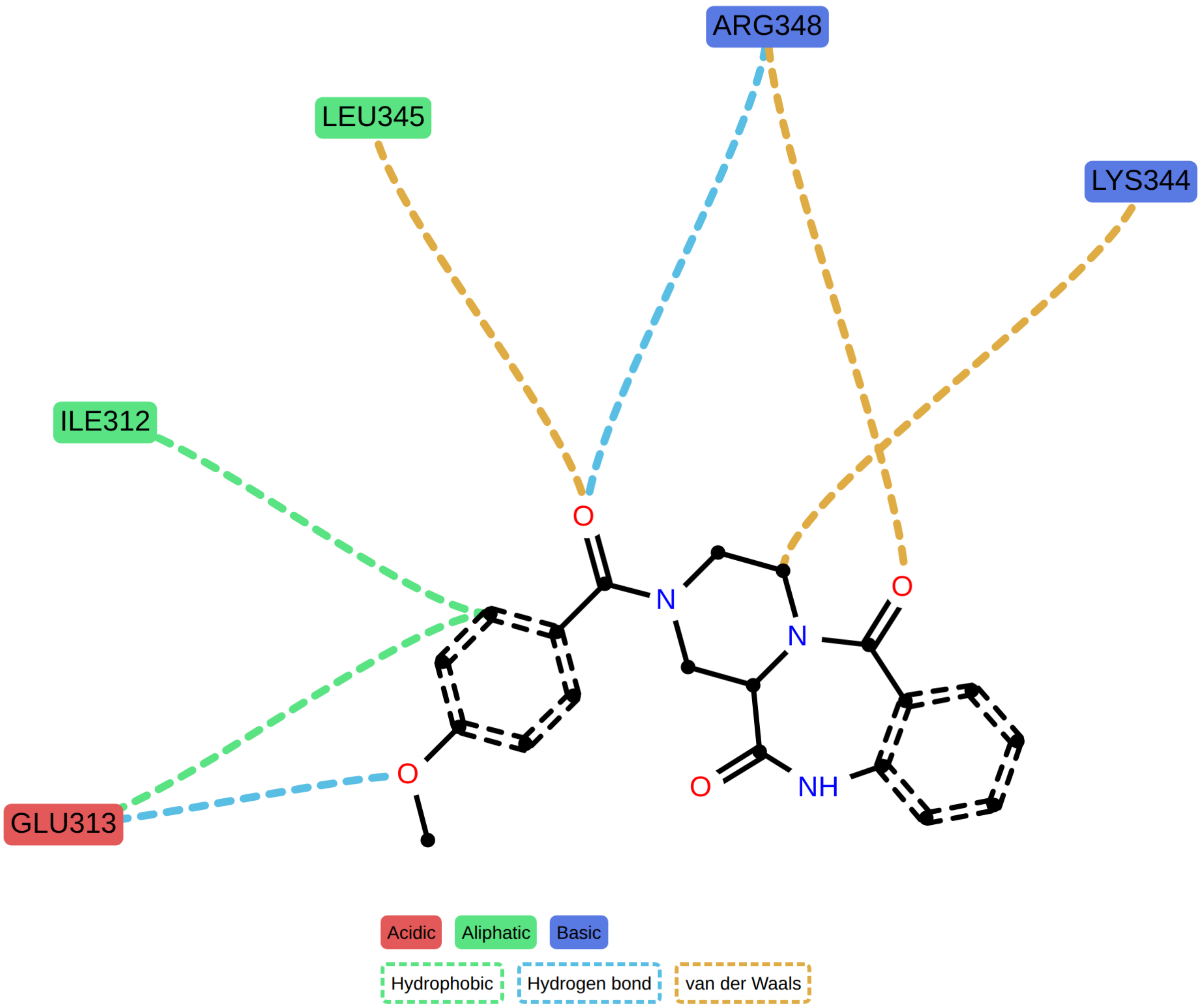


**Figure S5.** Two-dimensional interaction diagram of the top-ranked docking lead (CNP0062561.1; rank 1 of 24 by composite score, Supplementary Table S3) in the CHN1 interface-contact site, generated with ProLIF (LigNetwork). Residues are colored by type and edges by interaction type. The lead engages the Lys344 interface-contact region, forming hydrogen bonds to Glu313 and Arg348, a hydrophobic contact with Ile312, and van der Waals contacts with Lys344 and Leu345.

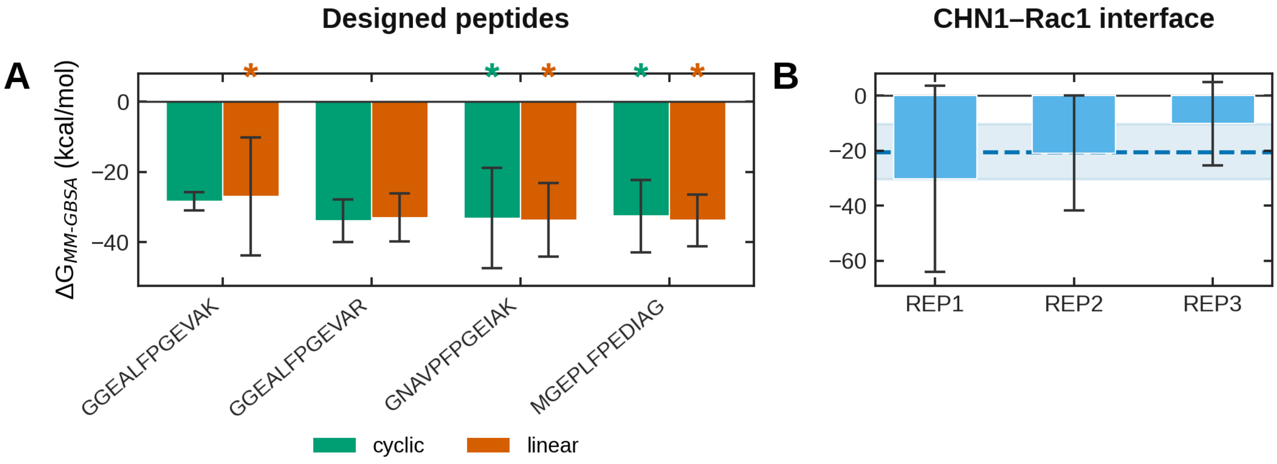


**Figure S6.** MM-GBSA end-point binding energies (single-trajectory, igb = 2). **(A)** Designed CHN1-directed peptides, cyclic (green) versus linear (orange). **(B)** CHN1–Rac1 interface across three independent 200 ns replicates; dashed line and band, inter-replicate mean ± SD. An asterisk marks a system evaluated from one trajectory, whose error bar is a per-frame.

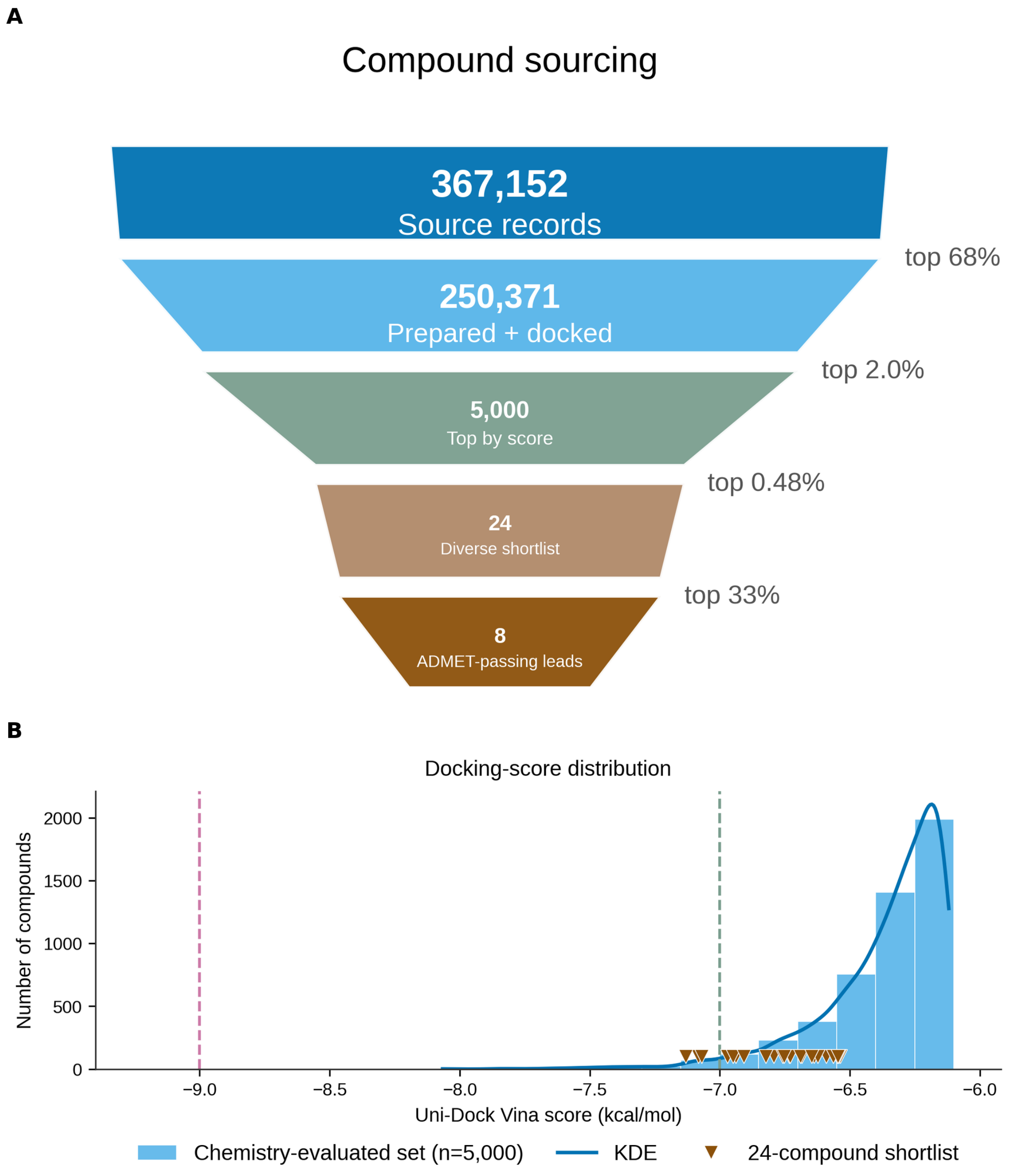


**Figure S7.** Essential dynamics of the lead CHN1–ligand complex (L13 = CNP0546584.1; 100 ns). **(A)** Projection of the CHN1 Cα motion onto the first two principal components, colored by simulation time; the first two components carry 80 % of the total positional variance. **(B)** Gibbs free-energy landscape over PC1 and PC2 (−RT ln P at 310 K), showing a single dominant basin, consistent with the ligand settling into one MD-refined binding mode.

### Natural-product leads

#### CNP0062561.1
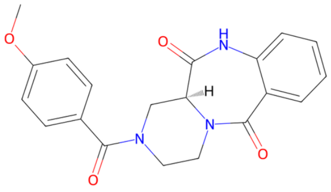

#### CNP0402905.1
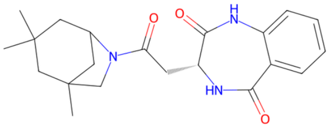

#### CNP0380192.1
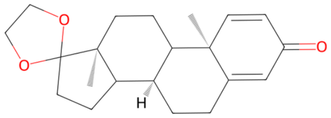

#### CNP0467273.1
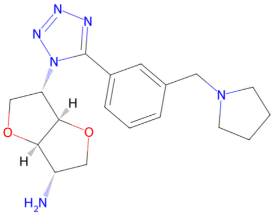

#### CNP0354493.0
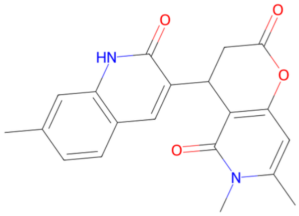

#### CNP0325351.4
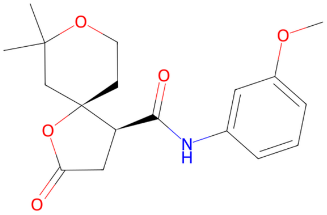

### CReM-optimized analogues

#### CHN1P_003575
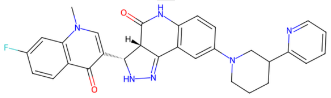

#### CHN1P_000746
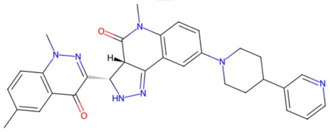

#### CHN1P_003652
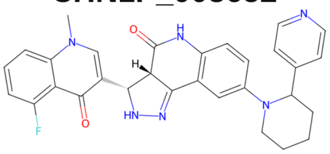

#### CHN1P_005034
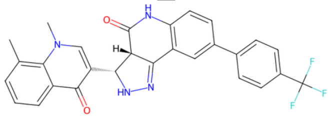

#### CHN1P_001140
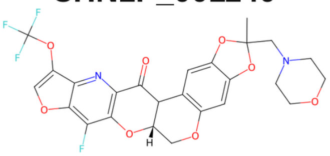

#### CHN1B_000442
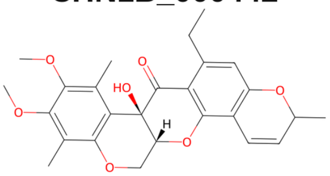

**Figure S8.** Full chemical panel from which the molecular-dynamics compounds were drawn: the eight Tier-1 COCONUT natural-product leads (top) and the eight top CReM-optimized analogues (bottom), each labeled with its compound ID. Docking scores and QED values are tabulated in the deposited shortlist. The four compounds carried into molecular dynamics (L01, L02, L13, L14) are shown in Fig. 4A.

### Supplementary Tables

**Table S1. Reported CHN1 substitutions, with domain and originating report.**

| Substitution | HGVS | Domain/region | Reference |
|---|---|---|---|
| L20F | p.Leu20Phe | alpha2-specific N-terminal/SH2 region | Miyake et al., 2008 |
| Y21C | p.Tyr21Cys | alpha2-specific N-terminal/SH2 region | Jurgens et al., 2025 |
| I126M | p.Ile126Met | alpha2-specific linker region | Miyake et al., 2008 |
| P141L | p.Pro141Leu | SH2-C1 linker | Chan et al., 2011 |
| Y143H | p.Tyr143His | SH2-C1 linker | Miyake et al., 2008 |
| Y148F | p.Tyr148Phe | SH2-C1 linker | Miyake et al., 2011 |
| F213L | p.Phe213Leu | C1 domain | Zhang et al., 2024 |
| F213V | p.Phe213Val | C1 domain | Zhou et al., 2020 |
| H217R | p.His217Arg | C1 domain | Zhang et al., 2024 |
| Y221H | p.Tyr221His | C1 domain | Angelini et al., 2021 |
| A223V | p.Ala223Val | C1 domain | Miyake et al., 2008 |
| N224S | p.Asn224Ser | C1 domain | Biler et al., 2017 |
| G228S | p.Gly228Ser | C1 domain/DAG-binding site | Miyake et al., 2008 |
| P252Q | p.Pro252Gln | C1 domain | Miyake et al., 2008 |
| P252S | p.Pro252Ser | C1 domain | Chan et al., 2011 |
| E313K | p.Glu313Lys | RacGAP domain/proximal Rac-binding site | Miyake et al., 2008 |

All substitutions were reported in patients with Duane retraction syndrome or a related ocular congenital cranial dysinnervation disorder. Tyr21Cys is classified as a variant of uncertain significance by its originating report and is listed for completeness.

**Table S2. Top-20 CHN1 interface residues by mean interface-contact score over three independent 200 ns CHN1–Rac1 replicates (mean ± inter-replicate SD).**

| CHN1 residue | Interface-contact score | Contact freq. | # Rac1 partners | Mean min dist (Å) | Salt-bridge freq. | H-bond freq. | n |
|---|---|---|---|---|---|---|---|
| LEU438 | 70.3 ± 10.1 | 0.99 ± 0.01 | 3.7 ± 0.6 | 3.73 ± 0.08 | 0.00 ± 0.00 | 0.00 ± 0.00 | 3 |
| ARG348 | 67.4 ± 10.2 | 0.96 ± 0.06 | 2.3 ± 0.6 | 3.33 ± 0.25 | 0.51 ± 0.19 | 0.28 ± 0.24 | 3 |
| PHE308 | 67.3 ± 20.1 | 0.93 ± 0.12 | 3.0 ± 1.7 | 3.61 ± 0.24 | 0.00 ± 0.00 | 0.00 ± 0.00 | 3 |
| ARG445 | 67.1 ± 12.4 | 0.95 ± 0.02 | 3.3 ± 1.2 | 3.32 ± 0.24 | 0.00 ± 0.00 | 0.35 ± 0.26 | 3 |
| ALA434 | 64.0 ± 12.6 | 0.96 ± 0.04 | 2.7 ± 1.5 | 3.59 ± 0.12 | 0.00 ± 0.00 | 0.00 ± 0.00 | 3 |
| PRO424 | 63.8 ± 5.9 | 1.00 ± 0.01 | 2.3 ± 0.6 | 3.55 ± 0.07 | 0.00 ± 0.00 | 0.01 ± 0.01 | 3 |
| ILE441 | 59.7 ± 11.1 | 0.93 ± 0.07 | 2.7 ± 0.6 | 3.83 ± 0.14 | 0.00 ± 0.00 | 0.00 ± 0.00 | 3 |
| ILE420 | 56.0 ± 25.9 | 0.77 ± 0.24 | 2.7 ± 2.1 | 3.41 ± 0.53 | 0.00 ± 0.00 | 0.37 ± 0.48 | 3 |
| LYS344 | 54.4 ± 33.5 | 0.72 ± 0.47 | 1.0 ± 1.0 | 3.00 ± 0.52 | 0.69 ± 0.51 | 0.67 ± 0.52 | 3 |
| MET435 | 51.7 ± 6.1 | 0.88 ± 0.08 | 1.7 ± 0.6 | 3.86 ± 0.06 | 0.00 ± 0.00 | 0.00 ± 0.00 | 3 |
| SER306 | 49.5 ± 37.2 | 0.64 ± 0.35 | 2.7 ± 3.8 | 3.43 ± 0.63 | 0.00 ± 0.00 | 0.37 ± 0.54 | 3 |
| LYS158 | 49.3 ± 21.1 | 0.54 ± 0.37 | 2.3 ± 1.5 | 2.94 ± 0.09 | 0.37 ± 0.44 | 0.46 ± 0.34 | 3 |
| ASP310 | 48.8 ± 20.0 | 0.76 ± 0.21 | 1.3 ± 0.6 | 3.54 ± 0.58 | 0.34 ± 0.48 | 0.33 ± 0.48 | 3 |
| SER309 | 48.3 ± 32.7 | 0.68 ± 0.45 | 2.0 ± 1.7 | 3.42 ± 0.49 | 0.00 ± 0.00 | 0.40 ± 0.52 | 3 |
| THR425 | 48.2 ± 19.4 | 0.82 ± 0.29 | 1.0 ± 0.0 | 3.49 ± 0.61 | 0.00 ± 0.00 | 0.35 ± 0.51 | 3 |
| GLU300 | 44.1 ± 29.6 | 0.52 ± 0.45 | 0.7 ± 0.6 | 2.71 ± 0.15 | 0.50 ± 0.43 | 0.49 ± 0.42 | 3 |
| VAL421 | 44.0 ± 12.1 | 0.77 ± 0.30 | 1.7 ± 0.6 | 3.81 ± 0.19 | 0.00 ± 0.00 | 0.01 ± 0.01 | 3 |
| GLU334 | 41.6 ± 17.2 | 0.52 ± 0.25 | 1.0 ± 1.0 | 3.05 ± 0.34 | 0.39 ± 0.22 | 0.37 ± 0.22 | 3 |
| ARG428 | 40.1 ± 21.8 | 0.57 ± 0.22 | 1.3 ± 1.2 | 3.63 ± 0.32 | 0.20 ± 0.22 | 0.18 ± 0.20 | 3 |
| GLY307 | 39.3 ± 29.7 | 0.63 ± 0.48 | 1.3 ± 1.5 | 3.61 ± 0.51 | 0.00 ± 0.00 | 0.33 ± 0.52 | 3 |

### Table S3. MM-GBSA and MM-PBSA end-point binding energies for the CHN1–ligand complexes (mean ± SD).

| Ligand | ΔG MM-GBSA (kcal/mol) | ΔG MM-PBSA (kcal/mol) | n | SD type |
|---|---|---|---|---|
| L13 | -34.54 ± 3.08 | -24.48 ± 4.00 | 3 | inter-replicate |
| L01 | -24.20 ± 5.93 | -21.38 ± 5.73 | 3 | inter-replicate |
| L02 | -25.25 ± 5.75 | -17.48 ± 8.73 | 3 | inter-replicate |
| L14 | -18.99 ± 5.01 | -14.76 ± 6.71 | 1 | per-frame |

### Table S4. CHN1–Rac1 interface MM-GBSA end-point binding energy across three independent replicates (generalized Born, OBC model).

| Replicate | ΔG MM-GBSA (kcal/mol) | SD (kcal/mol) | SD type |
|---|---|---|---|
| REP1 | -30.25 | 33.88 | per-frame (within run) |
| REP2 | -20.89 | 20.87 | per-frame (within run) |
| REP3 | -10.22 | 15.22 | per-frame (within run) |
| MEAN | -20.45 | 10.02 | inter-replicate |

### Table S5. Designed CHN1-directed peptides: last-10 ns Cα RMSF and MM-GBSA end-point binding energy (generalized Born, OBC model).

| Sequence | Form | Mean Cα RMSF (Å) | Max Cα RMSF (Å) | Replicate mean (Å) | Replicate SD (Å) | ΔG MM-GBSA (kcal/mol) | ΔG SD (kcal/mol) | n | SD type |
|---|---|---|---|---|---|---|---|---|---|
| GGEALFPGEVAK | Cyclic | 1.31 | 1.6 | 1.02 | 0.04 | -28.41 | 2.57 | 3 | inter-replicate |
| GGEALFPGEVAK | Linear | 2.17 | 4.33 | | | -27.05 | 16.78 | 1 | per-frame |
| GGEALFPGEVAR | Cyclic | 1.52 | 2.12 | 0.95 | 0.02 | -33.87 | 6.07 | 3 | inter-replicate |
| GGEALFPGEVAR | Linear | 1.14 | 1.43 | 1.28 | 0.11 | -33.02 | 6.85 | 3 | inter-replicate |
| GNAVPFPGEIAK | Cyclic | 1.88 | 3.23 | 1.57 | | -33.17 | 14.3 | 1 | per-frame |
| GNAVPFPGEIAK | Linear | 1.74 | 1.99 | | | -33.68 | 10.48 | 1 | per-frame |
| MGEPLFPEDIAG | Cyclic | 2.74 | 4.66 | 1.26 | | -32.62 | 10.27 | 1 | per-frame |
| MGEPLFPEDIAG | Linear | 1.98 | 3.1 | | | -33.84 | 7.38 | 1 | per-frame |

### Table S6. AlphaFold3 interactor screen ranked by the domain-normalized ipSAE (d0dom) (top 15 of 31 scored complexes).

| Rank | Interactor | Accession | ipSAE | ipSAE (d0dom) | AF3 ipTM | AF3 pTM | Confident contacts | Confidence class |
|---|---|---|---|---|---|---|---|---|
| 1 | NCK2 | O43639 | 0.675 | 0.789 | 0.57 | 0.47 | 104 | Strong |
| 2 | NCK1 | P16333 | 0.677 | 0.776 | 0.55 | 0.46 | 98 | Strong |
| 3 | RAC1 | P63000 | 0.453 | 0.61 | 0.76 | 0.61 | 186 | Good |
| 4 | NADK2 | Q4G0N4 | 0.017 | 0.213 | 0.25 | 0.44 | 9 | Weak |
| 5 | ADORA2A | P29274 | 0.03 | 0.043 | 0.4 | 0.5 | 18 | Weak |
| 6 | RPS3A | P61247 | 0.012 | 0.012 | 0.41 | 0.49 | 6 | Weak |
| 7 | KRAS | P01116 | 0 | 0 | 0.29 | 0.46 | 0 | Weak |
| 8 | SLC25A31 | Q9H0C2 | 0 | 0 | 0.15 | 0.41 | 0 | Weak |
| 9 | RNF123 | Q5XPI4 | 0 | 0 | 0.22 | 0.61 | 0 | Weak |
| 10 | PARK2 | O60260 | 0 | 0 | 0.18 | 0.34 | 0 | Weak |
| 11 | NIF3L1 | Q9GZT8 | 0 | 0 | 0.13 | 0.43 | 0 | Weak |
| 12 | NEF | P04601 | 0 | 0 | 0.17 | 0.46 | 0 | Weak |
| 13 | NAA20 | P61599 | 0 | 0 | 0.11 | 0.41 | 0 | Weak |
| 14 | MAPK8IP2 | Q13387 | 0 | 0 | 0.22 | 0.35 | 0 | Weak |
| 15 | MAPK1 | P28482 | 0 | 0 | 0.23 | 0.46 | 0 | Weak |

### Table S7. Trajectory-averaged molecular-dynamics parameters and DSSP secondary-structure content (mean ± inter-replicate SD).

| System | Backbone RMSD (Å) | CHN1 Cα RMSF (Å) | Rg (Å) | SASA ($nm^2$) | CHN1 intramol. H-bonds | n | α-helix | $3_{10}$-helix | β-strand | Turns | Bends | PP-II | Loops |
|---|---|---|---|---|---|---|---|---|---|---|---|---|---|
| apo CHN1 | 8.9 ± 0.9 | 2.4 ± 0.1 | 27.1 ± 0.3 | 248 ± 2 | 296 ± 3 | 3 | 172 ± 5 | 10 ± 0 | 38 ± 2 | 56 ± 3 | 56 ± 1 | 9 ± 1 | 98 ± 3 |
| CHN1–Rac1 complex | 8.7 ± 1.4 | 2.9 ± 0.4 | 30.0 ± 0.4 | 342 ± 2 | 296 ± 2 | 3 | 176 ± 1 | 9 ± 1 | 37 ± 2 | 53 ± 2 | 57 ± 1 | 9 ± 1 | 97 ± 2 |
| CHN1–ligand (CNP0546584.1) | 9.1 ± 1.0 | 3.0 ± 0.4 | 27.6 ± 0.4 | 250 ± 1 | 322 ± 2 | 3 | 179 ± 1 | 14 ± 2 | 37 ± 3 | 51 ± 1 | 56 ± 1 | 10 ± 2 | 90 ± 1 |

**Table S8. Composition of the 16-compound molecular-dynamics panel, mapping each L-number to its compound.**

| Label | Compound ID | Origin | Scaffold | Vina (kcal/mol) | QED |
|---|---|---|---|---|---|
| L01 | CHN1P_003575 | potency-optimized (CReM) | CHN1A_000536 | -9.797 | 0.423 |
| L02 | CHN1P_000746 | potency-optimized (CReM) | CHN1A_000304 | -9.564 | 0.432 |
| L03 | CHN1P_003652 | potency-optimized (CReM) | CHN1A_000506 | -9.467 | 0.416 |
| L04 | CHN1P_005034 | potency-optimized (CReM) | CHN1A_000355 | -9.391 | 0.398 |
| L05 | CHN1P_001140 | potency-optimized (CReM) | CHN1A_002387 | -9.372 | 0.446 |
| L06 | CHN1B_000442_f2e66051cf | drug-likeness-optimized (CReM) | CHN1Q_000183 | -8.212 | 0.756 |
| L07 | CHN1B_000648_ef235b4608 | drug-likeness-optimized (CReM) | CHN1T_000025 | -8.105 | 0.71 |
| L08 | CHN1B_000416_3559acb8ba | drug-likeness-optimized (CReM) | CHN1Q_000009 | -8.073 | 0.711 |
| L09 | CHN1B_000445_ad8e78773a | drug-likeness-optimized (CReM) | CHN1Q_000166 | -8.067 | 0.702 |
| L10 | CHN1B_000493_e97f394761 | drug-likeness-optimized (CReM) | CHN1Q_000029 | -8.007 | 0.702 |
| L11 | CNP0206284.2 | natural product (COCONUT) | Steroids | -6.939 | 0.698 |
| L12 | CNP0246271.0 | natural product (COCONUT) | — | -6.922 | 0.526 |
| L13 | CNP0546584.1 | natural product (COCONUT) | Nicotinic acid alkaloids | -6.826 | 0.554 |
| L14 | CNP0445170.1 | natural product (COCONUT) | — | -6.772 | 0.664 |
| L15 | CNP0124747.1 | natural product (COCONUT) | Triterpenoids | -6.729 | 0.73 |
| L16 | CNP0311700.4 | natural product (COCONUT) | Ornithine alkaloids | -6.713 | 0.654 |
